# A neural circuit for olfactory motion detection

**DOI:** 10.64898/2026.07.13.738327

**Authors:** Gustavo M. Santana, Harsh Vashistha, Thierry Emonet, Damon A. Clark

## Abstract

Movement is a defining feature of dynamic environments and a key variable encoded by sensory systems. In vision, neural mechanisms that extract motion direction have been studied extensively across species, revealing circuits that compare signals across space and time. Odor plumes likewise carry motion information in their spatiotemporal structure, which walking fruit flies detect and exploit to improve navigation (Brudner et al., 2025; Kadakia et al., 2022), independent of wind sensing. However, how olfactory circuits compute odor motion remains unknown. Here we identify circuitry in the *Drosophila* antennal lobe that transforms bilateral odor inputs into direction-selective signals that are propagated to higher brain regions. These direction-selective signals were widespread across most measured olfactory channels, spanning pheromones, food odors, and aversive cues. Directional responses were elicited by odor traces with naturalistic temporal statistics and tuned to inter-antennal delays of ≈40 ms. Connectomic, physiological, and behav-ioral analyses identified one GABAergic inhibitory neuron in the antennal lobe as a key mediator of direction selectivity. Measurements of dynamics in direction-selective neurons revealed ipsilateral excitation and delayed contralateral inhibition. In a minimal data-driven model, this mechanism was sufficient to generate the observed direction-selective responses. These results establish motion detection as a circuit-level olfactory computation and reveal how an early sensory circuit transforms spatiotemporal chemical signals into a neural representation of odor motion to guide navigation.

## MAIN

Objects moving through space generate characteristic spatiotemporal patterns that can be used to infer motion direction. A minimal motion detector requires separated sensors, a delay in processing between the sensors, and a nonlinear comparison of sensor output over time (Adelson and Bergen, 1985; Barlow and Levick, 1965; Hassenstein and Reichardt, 1956). In vision, direction-selective signals and the circuits that produce them are canonical examples of neural computation (Clark and Fitzgerald, 2024; Wei, 2018; Yang and Clandinin, 2018).

A circuit to detect olfactory motion must solve a similar problem to visual motion detection, but in a different sensory modality. Odor plumes are often intermittent, so an animal may encounter sequences of odor filaments rather than continuous spatial patterns (Murlis et al., 1992). Walking *Drosophila* use bilateral antennal sensing to respond to the direction and speed of odor motion, independently of wind sensing (Kadakia et al., 2022). While this behavior demonstrates that flies can infer the direction of odor motion by integrating odor signals from their two antennae, the neurons and circuit mechanisms that perform this computation are unknown. Here, we report the discovery of a direction-selective olfactory circuit in the *Drosophila* brain and characterize its properties.

### Odor motion drives direction-selective signals in the lateral horn

We first asked whether odor motion direction is represented in the lateral horn (Figure 1a), a region implicated in innate odor-guided behavior. We performed two-photon calcium imaging in immobilized flies while translating an apple-cider-vinegar odorized airflow back and forth across the fly’s head to generate a moving odor ribbon (Figure 1b-1e). Motion direction was defined relative to the imaged hemisphere: ipsi-first and contra-first motion directions were defined by whether the odor ribbon arrived first at the ipsilateral and contralateral antennae.

**Figure 1:**
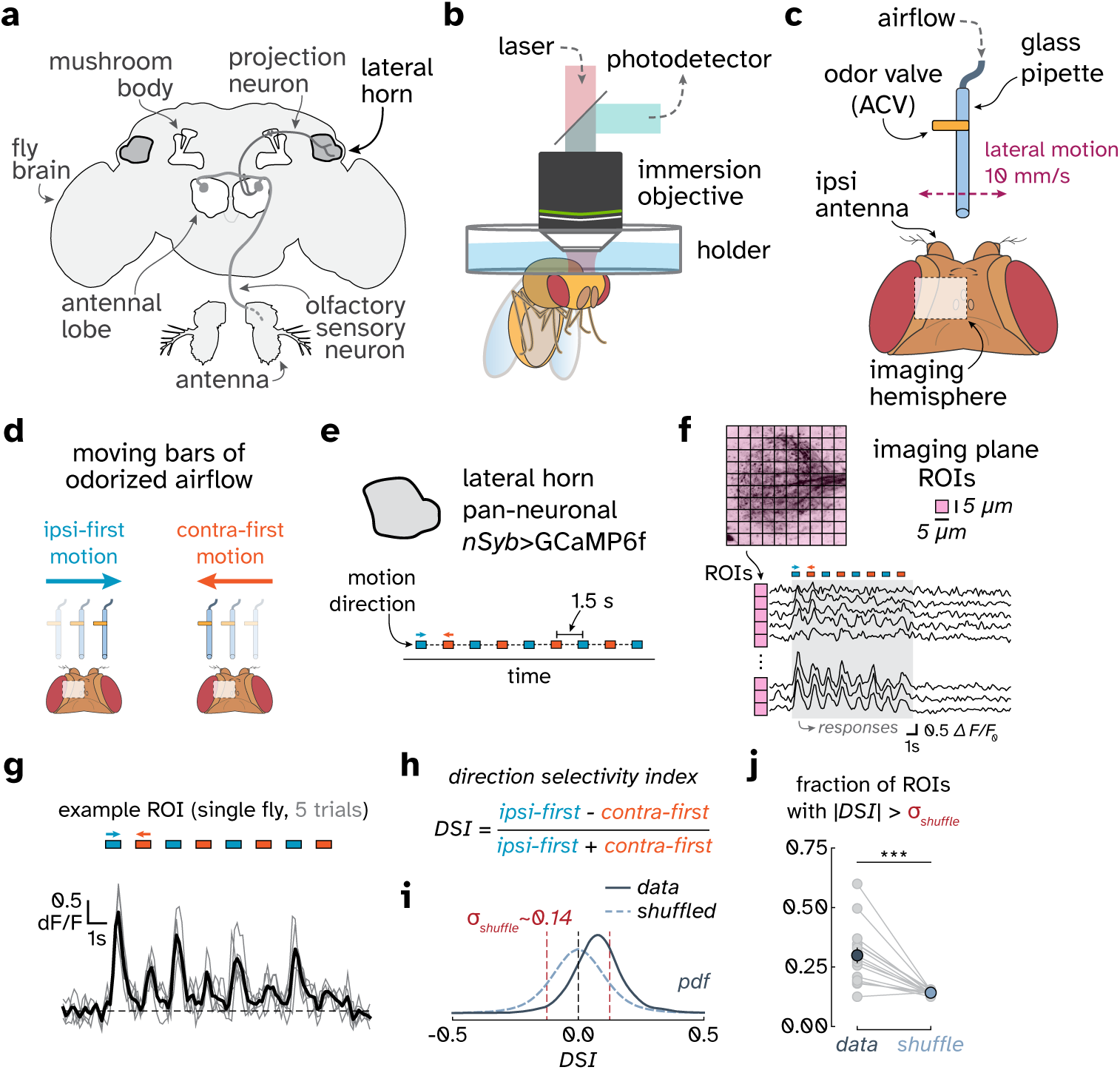
Moving odor ribbons evoke direction-selective activity in the lateral horn. **a.** Schematic of the adult fly olfactory pathway, showing olfactory receptor neurons, antennal lobe projection neurons, and their target regions, the mushroom body and the lateral horn. **b.** Two-photon imaging setup for recording calcium activity *in vivo* in tethered flies. **c.** Odor delivery apparatus. A glass pipette delivered undiluted apple-cider-vinegar in a narrow stream (1 L min*^−^*^1^) while translating laterally across the antennae at 10 mm s*^−^*^1^ (see Methods). The ipsilateral antenna was defined relative to the imaged hemisphere. **d.** Moving odorized ribbons generated ipsi-first or contra-first odor motion directions, defined by which antenna encountered the odorized airflow first. **e.** Stimulus trial structure for pan-neuronal lateral horn imaging (*nSyb*>GCaMP6f, see Methods). We presented flies with ipsi-first and contra-first moving odorized ribbons, alternating motion direction every ≈ 1.5 s. **f.** Grid-based definition of lateral horn regions of interest (ROIs; 5 *µ*m spacing) and example ROI calcium responses, shown as fractional fluorescence change (*dF* /*F* ). Responsive ROIs were retained using the inter-trial reliability criterion described in Methods. **g.** Example ROI response across five trials of alternating moving odor bars. Gray traces show individual trials; black trace shows the mean response. **h.** A direction selectivity index (DSI) was used to identify direction-selective ROIs. Each internal response peak was compared with the mean of its two neighboring peaks, which represented the opposite motion direction. Peak-wise contrasts were sign-coded so that positive values consistently indicated larger ipsi-first responses, and ROI DSI was computed as their mean (see Extended Data Fig. 1). **i.** DSI distribution across lateral horn ROIs from pan neuronal recordings and stimulus-label-shuffled controls. Vertical dashed red lines mark ± 1 s.d. of the shuffled distribution; n = 15. **j.** Fraction of lateral horn ROIs with |DSI| *> σ*_shuffle_ for real and shuffled stimulus labels. Each gray line denotes one fly; circles show mean ± s.e.m.; n = 15 flies; exact two-sided paired sign test, *P < 0.001*.

We imaged GCaMP6f (Chen et al., 2013) fluorescence in all neurons in the lateral horn (LH) and segmented them into regions of interest (ROIs). We found that calcium activity followed each pass of the moving odor ribbon (Figure 1f-1g). In most ROIs, the measured traces showed stronger responses to the ipsi-first direction than to the contra-first direction (Figure 1g), making ipsi-first the preferred direction and contra-first the null direction. To test whether this direction selectivity was above chance, we computed for each ROI a direction selectivity index (DSI), a normalized measure comparing mean responses to ipsi-first and contra-first motion (Extended Data Fig. 1). We then examined the distribution of DSI values across all ROIs in a fly. Many ROIs preferred ipsi-first motion, resulting in a modest but significant shift of the DSI distribution relative to a shuffled-label control (Figure 1i). Across flies, significantly more ROIs exceeded the shuffled threshold during odor motion than after label shuffling (Figure 1j). Thus, the LH contains neural representations of odor motion, with ipsi-first motion as the predominant preferred direction.

### Bilateral optogenetics isolates olfactory motion

Air-driven odor delivery couples odor timing to airflow and antennal mechanosensation. To isolate odor mo-tion from mechanosensation, we ceased using wind and instead expressed CsChrimson (Klapoetke et al., 2014) in all olfactory receptor neurons (ORNs). We used fiber-optic probes to independently activate these ORNs in the two antennae (Figure 2a-2b). Unilateral antennal stimulation produced stronger LH responses to ipsilateral stimulation (Figure 2c-2e), consistent with lateralized olfactory signals in the early olfactory system (Gaudry et al., 2013; Taisz et al., 2023).

**Figure 2:**
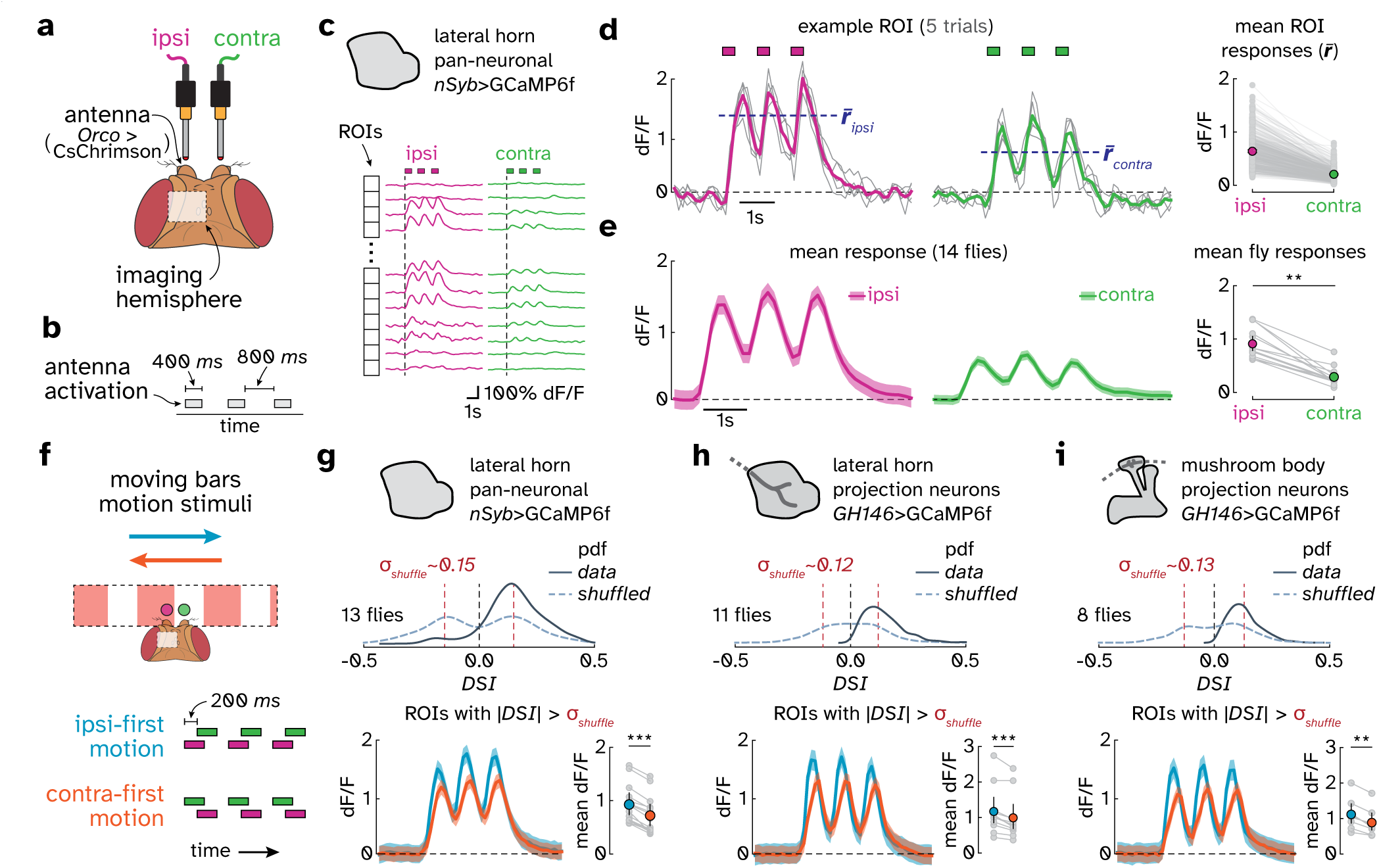
Bilateral optogenetic stimulation isolates olfactory motion and reveals projection neuron output signals. **a.** Fiber-optic probes for independent ipsilateral and contralateral stimulation of ORNs (*Orco*>CsChrimson, see Methods). Ipsi- and contralateral sides are defined relative to the imaged hemisphere. **b.** Unilateral optogenetic stimulus. Three 400 ms light pulses were delivered to either the ipsi or contra antenna, with 800 ms between pulses. **c.** Pan-neuronal lateral horn recordings using the same grid-based ROI definition as in Figure 1. **d.** Single-trial and mean dF/F responses to ipsilateral (left) and contralateral (middle) antenna activation in one ROI. Right, paired mean ROI responses for one fly across ROIs. n = 407 ROIs. **e.** Pan-neuronal lateral horn responses across flies. Left, mean response traces. Right, paired mean responses for individual flies. n = 14 flies; exact two-sided paired sign test, *P < 0.01*. **f.** Optogenetic moving bars as directional odor motion stimuli. Motion was generated by sequentially activating ORNs in each antenna with a fixed inter-antenna delay (Δ*_delay_* = 200 ms). **g.** Pan-neuronal lateral horn direction selectivity during moving bars. DSI was computed for each ROI as the normalized response difference between ipsi-first and contra-first motion (see Methods). Middle, DSI distribution for real and shuffled stimulus labels; dashed lines mark ± 1 s.d. (*σ*_shuffle_) of the shuffled distribution. Bottom, mean responses of |DSI| *> σ*_shuffle_ ROIs and fly-level response differences. n = 13 flies; exact two-sided paired sign test, *P < 0.001*. **h.** As in **g** but for antennal lobe projection neuron axon terminals in the lateral horn (GH146>GCaMP6f, see Methods for full genotype). n = 11 flies; exact two-sided paired sign test, *P < 0.001*. **i.** As in **h** but for antennal lobe projection neuron axon terminals in the mushroom body calyx. n = 8 flies; exact two-sided paired sign test, *P < 0.01*.

We then presented bilateral optogenetic odor motion sequences that simulated bars of odor that move across the antennae (Figure 2f). The ipsi-first and contra-first moving bar sequences differ only in the order of antennal activation. These matched sequences produced direction-selective LH responses with the same ipsi-first bias observed during air-driven odor motion (Figure 2g). Thus, differences in the timing of activity between the two antennae are sufficient to drive direction-selective LH activity.

### Projection neurons broadcast motion direction to higher olfactory centers

The LH receives abundant input from antennal lobe projection neurons (ALPNs). We therefore asked whether ALPN inputs already carried odor motion direction. To answer this question, we imaged axon terminals of a large subset of ALPNs in the LH in response to moving bars. These terminals also showed direction-selective responses with an ipsi-first preference (Figure 2h).

Direction selectivity in LH ALPN terminals could, in principle, reflect directional signals computed in the LH that influence ALPN terminal calcium signals there. To test whether ALPN terminals in other brain regions also showed direction-selective signals, we imaged ALPN terminals in the mushroom body calyx, a separate projection target. We found that calyx ALPN terminals also showed direction-selective responses with the same preferred direction (Figure 2i). These data suggest that the second-order ALPN olfactory neu-rons broadcast odor motion signals to multiple higher olfactory centers, but these data do not yet establish the site of motion computation.

### Direction selectivity emerges across antennal lobe glomeruli

To localize the motion computation, we compared ORN terminal and ALPN dendrite responses in many glomeruli across the antennal lobe to the same moving bars stimulus. Glomerular identity was assigned using odor landmarks, atlas position, and morphology (Benton et al., 2025; Knaden et al., 2012; Semmel-hack and Wang, 2009) (see Methods), yielding identified channels spanning pheromone, food, aversive, and mixed valence pathways (Figure 3a, Table 1).

**Figure 3:**
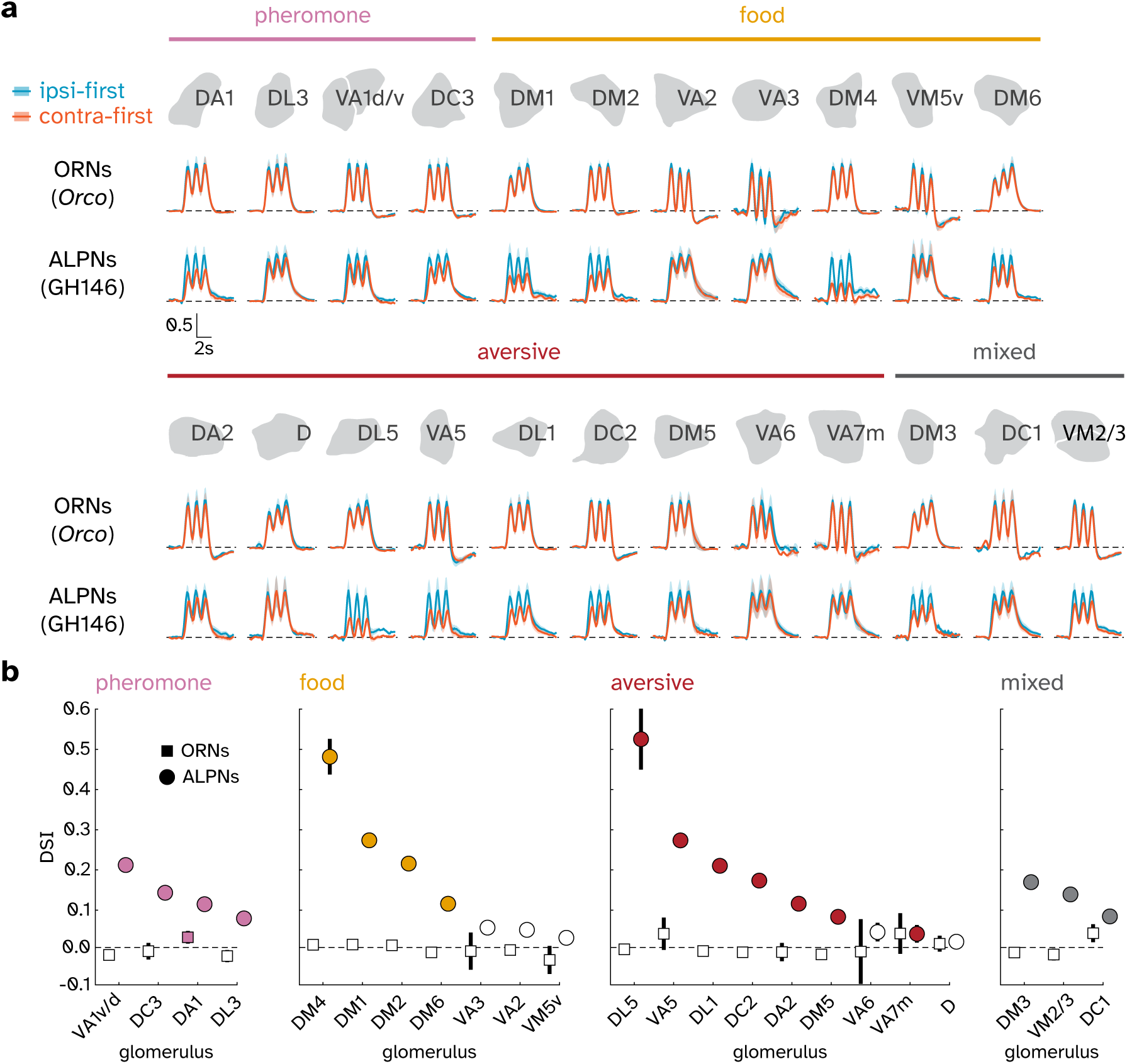
Projection neuron dendrites, but not ORN terminals, encode odor motion direction. **a.** Glomerular responses to ipsi-first and contra-first moving bars (as in Figure 2f) in ORN axon terminals (*Orco*>GCaMP6f) and ALPN dendrites (GH146>GCaMP6f). Columns show identified glomeruli grouped by ecological relevance: pheromones, foods, aversive, and mixed signals. See Table 1 for odorant and valence information for each glomerulus. Traces show normalized dF/F responses (mean ± s.e.m.), scaled by the mean peak response for each glomerular label; n = 5-8 flies for each glomerulus. **b.** Ranked glomerular DSI for the ORN terminal and ALPN dendrite responses shown in **a**, grouped by ecological relevance. DSI was computed as in Figure 2. Symbols show mean ± s.e.m.; error bars may be smaller than symbols. Filled symbols indicate DSI significantly different from zero; two-sided Wilcoxon signed-rank tests followed by Benjamini–Hochberg correction within each test family (q < 0.05, false-discovery-rate-adjusted *P* value); empty symbols indicate not significant.

**Table 1:** Direction selectivity index and odor annotations for glomeruli used in the direction-selectivity analysis.

| Glomerulus | DSI | Sensilla | Receptor | Odor valence / quality | Key odorants |
| --- | --- | --- | --- | --- | --- |
| **Pheromones** |  |  |  |  |  |
| **DA1** | 0.212 | at1 | Or67d | mating/aggression; pheromones | 11-cis-vaccenyl acetate |
| **DC3** | 0.143 | ai2 | Or83c | pheromones | farnesol |
| **VA1v** | 0.115 | at4 | Or47b | appetitive; pheromones | palmitoleic acid, methyl laurate |
| **VA1d** | 0.115 | at4 | Or88a | pheromones | methyl laurate, methyl myristate, methyl palmitate |
| **DL3** | 0.079 | at4 | Or65a/(Or65b/c) | pheromones | 11-cis-vaccenyl acetate (weak) |
| **Food** |  |  |  |  |  |
| **DM4** | 0.482 | ab2 | Or59b | appetitive; yeasty | methyl acetate, ethyl acetate, ketones |
| **DM1** | 0.274 | ab1 | Or42b | appetitive; yeasty | esters (e.g. ethyl acetate, ethyl propionate), 3-hexanone |
| **DM2** | 0.216 | ab3 | Or22a/(Or22b) | appetitive; alcoholic fermentation; fruity | esters (e.g. methyl hexanoate, ethyl hexanoate) |
| **VM2** | 0.139 | ab8 | Or43b | appetitive; fruity | esters (e.g. ethyl butyrate), alcohols (e.g. Z2-hexenol) |
| **DM6** | 0.116 | ab10 | Or67a | yeasty | esters (e.g. ethyl benzoate, pentyl acetate), 6-methyl-5-hepten-2-one, pentanoic acid |
| **VA3** | 0.057 | ab9 | Or67b | appetitive; yeasty; fruity | alcohols (e.g. trans-3-hexen-1-ol, 1-hexanol), acetophenone |
| **VA2** | 0.051 | ab1 | Or92a | appetitive; alcoholic fermentation | 2,3-butanedione |
| **VM5v** | 0.031 | ab7 | Or98a | appetitive; yeasty; fruity | esters (e.g. ethyl benzoate), alcohols (e.g. E2-hexenol), ketones (e.g. 6-methyl-5-hepten-2-one) |
| **Aversive** |  |  |  |  |  |
| **DL5** | 0.525 | ab4 | Or7a | aversive; alcoholic fermentation; dangerous; plant matter | E2-hexenal, E2-hexenol |
| **VA5** | 0.274 | ab11 | Or49b | aversive; animal matter | indoles, methylphenols |
| **DC2** | 0.174 | ab6 | Or13a | aversive; plant matter | alcohols (e.g. 1-octen-3-ol) |
| **DA2** | 0.116 | ab4 | Or56a/(Or33a) | aversive; dangerous | geosmin |
| **VA7m** | 0.041 | ab6 | Or46aB | aversive; dangerous | methylphenols |
| **D** | 0.021 | ab9 | Or69aA/B | aversive; dangerous | Z4-undecenal, carvone, terpenoid alcohols (e.g. linalool), esters |
| **Mixed** |  |  |  |  |  |
| **DL1** | 0.210 | ab1 | Or10a/(Gr10a) | aversive; fruity | methyl salicylate and other esters, acetophenone |
| **DM3** | 0.170 | ab5 | Or47a/(Or33b) | aversive; yeasty; fruity; dangerous | esters (e.g. pentyl acetate, methyl hexanoate), 3-methylthio-1-propanol |
| **VM3** | 0.139 | ab8 | Or9a | aversive; alcoholic fermentation | esters (e.g. ethyl-3-hydroxybutyrate), alcohols (e.g. 2-pentanol), ketones (e.g. 3-hydroxy-2-butanone) |
| **DC1** | 0.084 | ai3 | Or19a/(Or19b) | fruity; dangerous | terpenes (e.g. valencene, limonene) |
| **DM5** | 0.083 | ab2 | Or85a/(Or33b) | aversive; yeasty | ethyl-3-hydroxybutyrate |
| **VA6** | 0.045 | ab5 | Or82a | aversive; yeasty; plant matter | geranyl acetate |

ORN terminals responded robustly to moving bars but showed little difference between ipsi-first and contra-first motion across the imaged glomeruli (Figure 3a; Extended Data Fig. 3). ALPN dendrites, in con-trast, showed direction-selective responses in many of the same glomeruli, some with large differences be-tween preferred and null direction responses (Figure 3a). Matched ALPN ROIs in opposite hemispheres showed the corresponding reversal of motion direction preference, confirming that the signal was defined relative to antenna order (Extended Data Fig. 4). These measurements place the earliest computation of odor motion direction one synapse away from the periphery.

ALPN direction selectivity was broad but heterogeneous. Ranking ALPNs by the strength of their DSI revealed the strongest ipsi-first bias in DL5 (aversive) and DM4 (food), clear direction selectivity in DM1, VA5, DM2, DL1 and DC2, and weak or near-zero selectivity in VA7m, VM5v and D (Figure 3b). The ipsi-first preference was observed irrespective of whether the glomeruli responded to attractive or aversive odors. Thus, odor motion direction emerges as a distributed antennal lobe output feature with channel-specific strength and ipsi-first direction preference in each hemisphere.

### DA1 projection neurons resolve fast dynamics of odor motion

Because direction-selective output was present in both specialized and broadly tuned odor channels, we next selected DA1, a glomerulus responsive to the pheromone cVA (Kurtovic et al., 2007), as a pathway to characterize how dynamic antennal inputs are transformed into ALPN direction-selective signals.

Computing the direction of motion requires comparing inputs across relative delays (Borst and Egel-haaf, 1989). To measure the delay tuning of DA1 ALPNs to moving bars, we varied the inter-antenna delay of the stimulus (Figure 4a). DA1 ALPNs responded more strongly to ipsi-first than contra-first sequences at delays of over 100 ms (Figure 4b-4c), while DSI first became significant at ≈25 ms (Figure 4d). This func-tional dependency is expected from standard models of direction selectivity and is consistent with behav-ioral responses to olfactory motion (Kadakia et al., 2022).

**Figure 4:**
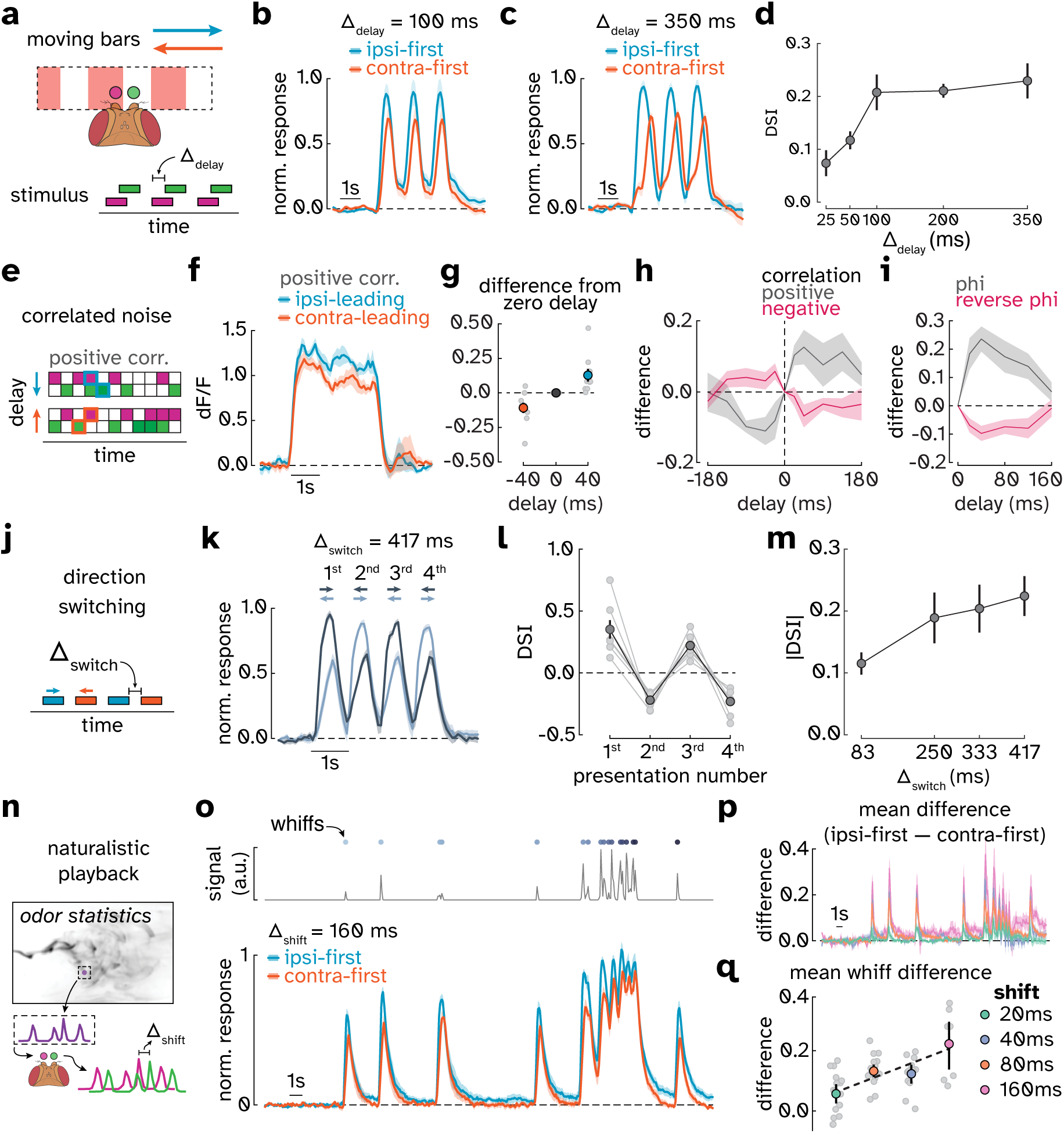
DA1 projection neurons resolve fast dynamics of odor motion. **a.** Moving bars stimuli, as in Figure 2f, was used to vary inter-antenna delay Δ_delay_ between ipsi- and contralateral ORN activation. **b.** Example DA1 ALPN responses to ipsi-first and contra-first moving bars at Δ_delay_ = 100 ms. Normalized response is dF/F normalized to the mean peak response across trials, per fly (see Methods for details). Traces show mean ± s.e.m.; n = 10 flies. **c.** As in **b** but for Δ_delay_ =350 ms; n = 14 flies. **d.** Mean DSI as a function of inter-antenna delay. Circles show mean ± s.e.m.; n = 10/14/10/11/14 for 25/50/100/200/350 ms; exact two-sided sign tests versus zero, *P = 0.021*, *P < 0.01*, *P < 0.01*, *P < 0.001*, and *P < 0.001*. **e.** Correlated-noise stimulus used to vary the imposed inter-antennal delay and correlation sign. Ipsilateral stimulus was binary white noise sampled at 50 Hz, and the contralateral stimulus was a delayed copy (correlation +1) or delayed complement (correlation -1). Contralateral-first stimuli swapped the two antenna assignments. Stimulus construction is described in Methods and shown in Extended Data Fig. 5. **f.** Time trace (mean ± s.e.m.) of DA1 responses to positively correlated noise when the ipsi- or contralateral antenna led by 40 ms. n = 8-9 flies. **g.** Response difference relative to the zero-delay condition as a function of imposed delay (-40, 0 and 40 ms) for positively correlated noise. Gray circles show individual flies; colored circles show mean ± s.e.m.; n = 8-9 flies. **h.** Response difference relative to the zero-delay condition for positively and negatively correlated noise as a function of imposed delay. Negative delays correspond to contralateral-leading stimuli (see Extended Data Fig. 6 for per-delay leading-versus-lagging comparisons). Traces show mean ± s.e.m.; n = 8-9 flies. **i.** For each delay, the net response to positive correlations (ipsi-leading − contra-leading; labeled phi) and the net response to negative correlations (ipsi-leading − contra-leading; labeled reverse-phi). Traces show mean ± s.e.m.; n = 8-9 flies. **j.** Direction-switching stimulus in which moving bars (as in **a**) were presented sequentially in alternating motion directions. Δ_switch_ is the interval from the offset of the trailing bar to the onset of the next bar. **k.** DA1 ALPN responses during repeated direction switches at Δ_switch_ = 417 ms. Dark trace begins with ipsi-first motion, and light trace begins with contra-first motion. Traces are mean ± s.e.m.; n = 7 flies. **l.** Signed DSI for successive bar presentations. DSI was computed as 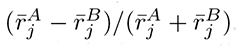, where *A* and *B* denote the traces that began with ipsi-first and contra-first motion, respectively. Because the traces exchanged motion direction at each presentation, the DSI sign alternated after each direction switch. Gray circles show individual flies; dark circles show mean ± s.e.m.; n = 7 flies; exact two-sided sign tests versus zero for each presentation, *P = 0.0156*. **m.** Mean absolute DSI across presentations 2–4 as a function of switch interval. Circles show mean ± s.e.m.; n = 7 flies; exact within-fly permutation trend test, *P < 0.0001*. **n.** Naturalistic optogenetic stimulus generated from a measured odor-plume waveform. An inter-antenna temporal shift Δ_shift_ was imposed between the two antennal input streams to simulate plume motion. **o.** Top: naturalistic stimulus waveform. Bottom: DA1 responses to ipsi-first and contra-first naturalistic stimuli at Δ_shift_ = 160 ms. Traces show mean ± s.e.m.; n = 7 flies. **p.** Mean whiff-aligned response difference traces (ipsi-first − contra-first) across imposed shifts. Traces show mean ± s.e.m.; n = 7-18 flies. **q.** Time-averaged whiff response difference as a function of imposed shift. Gray circles show individual flies; colored circles show mean ± s.e.m.; black dashed line shows descriptive least-squares fit (*r*2 = 0.856); n = 17/18/12/7 flies for 20/40/80/160 ms shifts; exact two-sided sign tests versus zero, *P = 0.049*, *P < 0.0001*, *P < 0.001*, and *P < 0.01*.

To better measure the tuning of this motion detector to different delays, we stimulated the antennae with pairwise correlations in intensity. Such correlations are a hallmark of the spatiotemporal patterns gen-erated by translational motion (Fitzgerald et al., 2011; Potters and Bialek, 1994) and central to canonical models of visual motion detection (Adelson and Bergen, 1985; Hassenstein and Reichardt, 1956). Our stim-uli consisted of binary stochastic signals with imposed correlations between the antennae at defined tem-poral offsets (Figure 4e; stimulus construction in Extended Data Fig. 5). These stimuli were adapted from visual motion stimuli whose perceived direction reverses when the intensity correlations are inverted (Adelson and Bergen, 1985). The positively correlated noise produced larger responses when the correlation was oriented in the preferred direction (Figure 4f-4g). The directional response varied systematically with imposed delay over the range from 20 to 120 ms, with peak responses at ≈40 ms (Figure 4h-4i; per-delay responses in Extended Data Fig. 6). The negatively correlated stimuli produced opposite results, in which the response was larger when the negative correlation was oriented in the null direction. Thus, this odor motion detector is sensitive to the sign of pairwise correlations, consistent with canonical models for motion detection (Adelson and Bergen, 1985; Hassenstein and Reichardt, 1956). The tuning curve describes an effective delay in the motion computation (Salazar-Gatzimas et al., 2016), consistent with our earlier measurements (Figure 4d). Moreover, responses to negative odor correlations provide a neural correlate of the behavioral sensitivity to negative correlations (Kadakia et al., 2022). It also represents an olfactory analogue of the visual reverse-phi motion illusion (Anstis, 1970; Hassenstein and Reichardt, 1956).

Consistent with a correlation-based motion detector (Egelhaaf et al., 1989; Reichardt and Guo, 1986), the DA1 ALPNs respond to drifting sinusoidal patterns with an amplitude that depends on the ratio of its velocity to its wavelength (Extended Data Fig. 7). This phenomenology is similar to visual motion detectors in the fly (Creamer et al., 2018; Haag et al., 2004), mammalian retina (Grzywacz and Amthor, 2007; He and Levick, 2000), and visual cortex (Holub and Morton-Gibson, 1981; Priebe et al., 2006).

Odor motion direction can change rapidly in odor plumes. To measure how quickly DA1 ALPNs could follow changing directional input, we presented moving bars that switched back and forth in direction (Figure 4j). The responses tracked these switches: ipsi-first bars produced larger responses than contra-first bars (Figure 4k-4l), even as the directional switches occurred with gaps of less than 100 ms (Figure 4m), showing that DA1 tracks rapid changes in direction.

Finally, we asked whether DA1 directional signals respond rapidly enough to track the transient fluctuations generated by individual odor filaments passing over a fly. To test this, we generated an intermittent optogenetic stimulus that simulates the arrival of brief odor bursts, or whiffs, transported by the wind (Gorur-Shandilya et al., 2017). The whiff concentrations and durations are power law-distributed, consistent with turbulent statistics (Celani et al., 2014). To simulate the movement of a plume across the two antennae, we imposed temporal shifts between the two antennal input streams (Figure 4n). DA1 whiff-locked responses depended on the relative whiff timing across the antennae: ipsi-leading whiffs consistently evoked larger responses than contra-leading whiffs (Figure 4o), and this difference increased with the imposed inter-antennal delay (Figure 4p-4q). These results show that DA1 direction selectivity is preserved for plume-like temporal signals and can encode motion generated by individual odor filaments.

Overall, this suite of experiments characterizes the motion computation in DA1 ALPNs and finds that it appears algorithmically similar to canonical visual computations of motion, while also being sensitive to naturalistic odor statistics.

### GABAergic inhibition is required for DA1 direction selectivity

DA1 response tuning suggested a simple motion detector with one delay and a nonlinearity (Figure 4f-4i, Extended Data Fig. 7). Because ALPN null-direction responses were suppressed relative to simultaneous bilateral pulses (Extended Data Fig. 2), we hypothesized that the mechanism might be analogous to the Barlow-Levick model of direction selectivity in the mammalian retina (Barlow and Levick, 1965). There, delayed inhibition from one sensor cancels non-delayed excitation from the other during null-direction motion (Figure 5a). As a broad test of this possibility, we recorded DA1 ALPN responses to moving bars after bath exchange with either fresh saline or application of GABA receptor antagonists (Figure 5b-5c). GABA receptor blockade significantly reduced response directionality toward zero across flies (Figure 5d), showing that inhibition is required for normal DA1 motion responses, consistent with a delayed inhibition mechanism.

**Figure 5:**
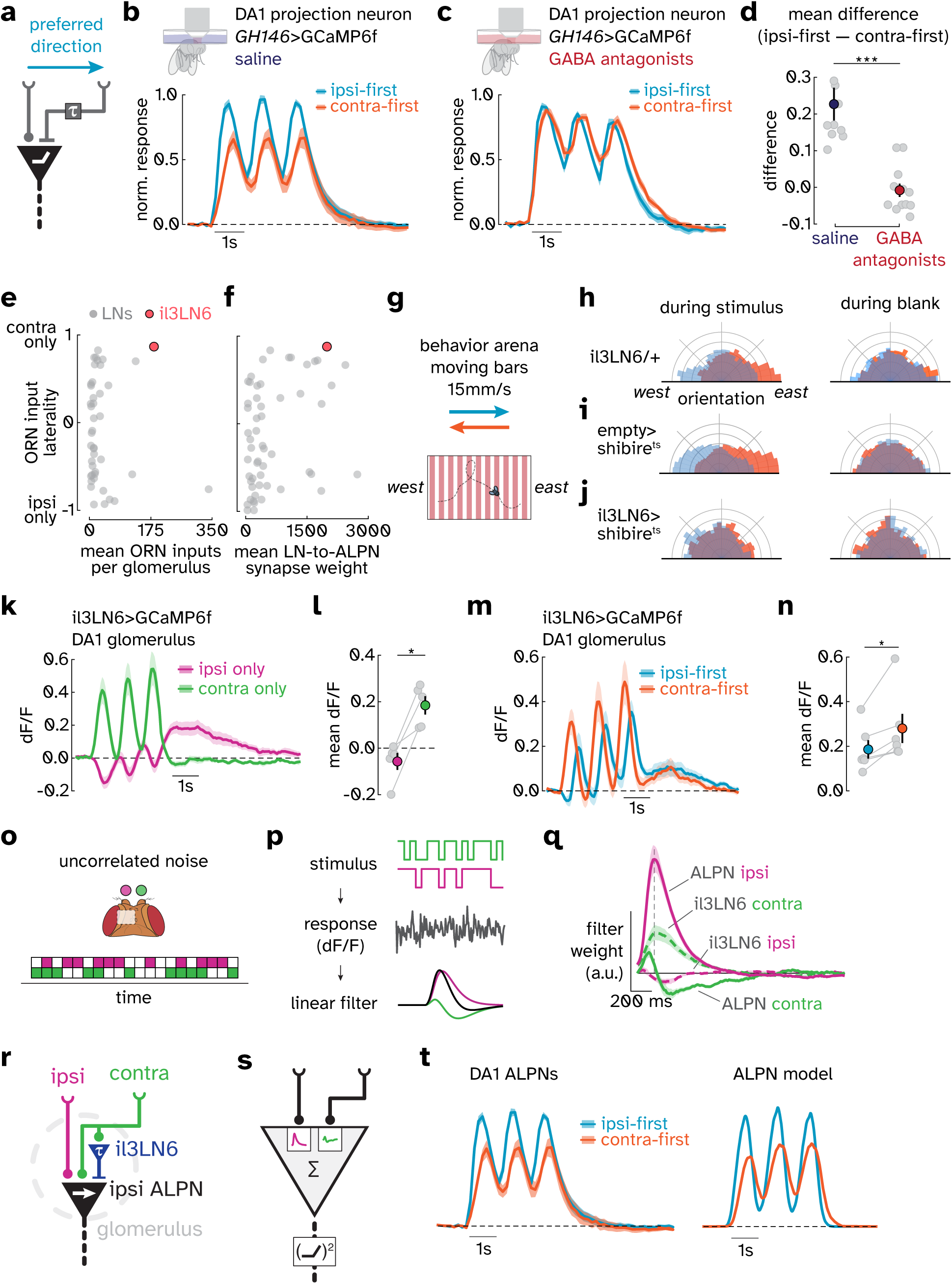
GABAergic inhibition and il3LN6 shape DA1 direction selectivity and behavior. **a.** Schematic of a simplified Barlow-Levick model of direction selectivity with delayed inhibition. In this diagram, the preferred direction is ipsi-first motion. **b.** Normalized DA1 ALPN responses to ipsi-first and contra-first moving bars after bath exchange with saline imaging solution. Traces show mean ± s.e.m.; n = 10 flies. **c.** Normalized DA1 ALPN responses after bath application of GABA receptor antagonists picrotoxin (20 µM) and CGP 54626 (50 µM). Traces show mean ± s.e.m.; n = 12 flies. **d.** Time-averaged response difference (ipsi-first − contra-first) in saline and GABA antagonist treated flies. Gray circles show individual flies; colored circles show mean ± s.e.m.; saline n = 10 flies; GABA antagonists n = 12 flies; two-sided Mann–Whitney U test, *P < 0.0002*. **e.** FlyWire connectome analysis of antennal lobe local neuron (LN) types. Each circle represents one LN type. The y axis shows mean ORN input laterality, computed as the normalized difference between contralateral and ipsilateral ORN inputs. The x axis shows mean ORN-to-LN synapse count per innervated glomerulus. il3LN6 is highlighted in red. **f.** Same ORN input laterality metric as in **e**, plotted against mean LN-to-ALPN synapse weight. il3LN6 is highlighted in red. **g.** Schematic of behavioral arena for measuring freely walking fly orientation while presenting optogenetic moving bars. **h.** Orientation distributions for il3LN6/+ genetic control flies during optogenetic moving bar stimulus presentation and blank periods (see Methods for silencing protocol). n = 286 paired trajectories. **i.** As in **h** but for empty>shibire^ts^ genetic control flies. n = 275 paired trajectories. **j.** As in **h** but for il3LN6-silenced (il3LN6>shibire^ts^) flies. n = 206 paired trajectories. **k.** il3LN6 responses in the DA1 glomerulus to unilateral ORN activation. Traces show mean ± s.e.m.; n = 5 flies. **l.** Time-average of responses in **k**. Gray lines show individual flies; colored circles show mean ± s.e.m.; n = 5 flies; one-sided exact Wilcoxon signed-rank test, *P = 0.03125*. **m.** il3LN6 responses in the DA1 glomerulus to ipsi-first and contra-first moving bars. Traces show mean ± s.e.m.; n = 6 flies. **n.** Time-average of responses in **m**. Gray lines show individual flies; colored circles show mean ± s.e.m.; n = 6 flies; two-sided exact Wilcoxon signed-rank test, *P = 0.03125*. **o.** Bilateral optogenetic noise stimulus used to estimate linear response filters. Each frame independently activated the ipsilateral antenna, contralateral antenna, both antennae, or neither antenna. **p.** Least-squares fitting workflow for estimating ipsilateral and contralateral response filters from measured calcium traces (see Methods). **q.** Estimated linear response filters for DA1 ALPNs and il3LN6 responses to ipsilateral and contralateral antennal input. Traces show mean ± s.e.m.; n = 14 DA1 ALPNs; n = 10 for il3LN6. **r.** Proposed direction selectivity circuit schematic. il3LN6 receives contralaterally biased ORN input and provides delayed inhibitory input to the DA1 projection neuron pathway. **s.** Minimal data-driven model schematic. Measured DA1 ALPN filters were convolved with the moving bar stimulus, summed, and passed through a squared rectifying nonlinearity (see Methods). **t.** Time-averaged *in vivo* DA1 ALPN responses (left) and model responses (right) to ipsi-first and contra-first moving bars.

### Connectomics identifies a candidate inhibitory pathway

The pharmacology identified inhibition as a key component but did not identify its origin. Antennal lobe local neurons are predominantly inhibitory but vary widely in morphology and connectivity (Chou et al., 2010; Liou et al., 2018; Schlegel et al., 2021). We therefore searched the antennal lobe connectomes (Dorkenwald et al., 2024; Scheffer et al., 2020; Schlegel et al., 2024) for local neurons (LNs) with two expected properties in each glomerulus: biased input from the contralateral compared to ipsilateral ORNs and substantial output onto ALPNs. These criteria should identify local neurons positioned to suppress ALPN responses during contra-first, null-direction motion. We plotted the fraction of ORN input from the ipsi- and contralateral antennae over all LNs and all glomeruli (Extended Data Fig. 8). Compared to other LNs, il3LN6 receives a very large fraction of its ORN inputs from the contralateral antenna and these connections are very strong (Figure 5e). Interestingly, this same neuron has been implicated in odor gradient detection (Taisz et al., 2023). Moreover, il3LN6 is also strongly connected to ALPNs (Figure 5f). This pattern of connectivity nominates il3LN6 as a strong candidate origin of lateralized inhibition for shaping ALPN direction selectivity.

### il3LN6 carries direction-selective signals and shapes behavior

If il3LN6 contributes to odor motion computation, silencing it should specifically affect behavioral responses to odor motion. We recorded freely-walking flies in an arena while presenting optogenetic moving bars and measured fly orientation relative to the bar motion direction (Figure 5g). Control flies reoriented against the motion of the bars, consistent with prior measurements (Kadakia et al., 2022), an effect that dis-appears during blank periods (Figure 5h-5i; Extended Data Fig. 9a-9b). In il3LN6-silenced flies, orientation against bar motion was strongly reduced, and there was no significant difference between the presentation and blank periods (Figure 5j; Extended Data Fig. 9c). The il3LN6-silenced fly behavior was significantly less oriented than either parental genetic control (Extended Data Fig. 9). These flies could still navigate an odor ribbon in wind (Extended Data Fig. 10), showing that silencing il3LN6 did not eliminate all odor detection or odor-guided behavior.

We next asked whether il3LN6 physiology matched the connectomic prediction. Unilateral optogenetic activation of ORNs in the contralateral antenna strongly excited il3LN6, whereas ipsilateral ORN activation produced weak suppression (Figure 5k-5l). In the mammalian retina, inhibitory amacrine cells that synapse onto direction-selective retinal ganglion cells are themselves direction-selective and provide stronger in-hibitory input when stimuli are directed in the null direction of the ganglion cells (Briggman et al., 2011; Eu-ler et al., 2002). Similarly, il3LN6 responded more strongly to contra-first than ipsi-first moving bar stimuli (Figure 5m-5n), the opposite preferred direction from downstream DA1 ALPNs.

### A minimal data-driven model qualitatively reproduces measured direction selectivity

The physiology suggests two non-exclusive mechanisms for direction selectivity in ALPNs: (1) il3LN6 itself carries a direction-selective inhibitory signal (Figure 5m-5n) that could generate ALPN direction selectiv-ity, but (2) il3LN6 could nonetheless also provide delayed inhibitory input to DA1 ALPNs to generate direc-tion selectivity. We therefore tested whether a delayed inhibition model could account for the directional measurements made in ALPNs. We first presented uncorrelated stimulus noise that independently acti-vated ORNs in the two antennae (Figure 5o-5p), allowing us to measure delays by estimating the impulse response filters of DA1 ALPNs and il3LN6 neurons to ipsilateral and contralateral antennal activation (Fig-ure 5q).

The measured filters captured the temporal weighting of ipsilateral and contralateral antennal inputs. Consistent with our prior results, il3LN6 was excited by contralateral input and modestly suppressed by ip-silateral inputs. The ALPN ipsilateral kernel showed strong, fast excitation. But the ALPN response to con-tralateral input showed an initial activation followed by suppression. These filters are consistent with fast excitation by ipsilateral ORNs followed by delayed contralateral inhibition from il3LN6 (Figure 5r). Interest-ingly, a similar computational structure emerges from optimizing a small neural network to make naviga-tional decisions in odor plumes (Brudner et al., 2025).

To assess whether these filters could explain direction selectivity in DA1 ALPNs, we built a simple data-driven model using the measured filters from the ALPN calcium signals. Ipsilateral and contralateral moving bar stimulus traces were each convolved with their corresponding filters, and the filtered responses were summed and passed through a squared rectifying nonlinearity (Figure 5s). This minimal model qualitatively reproduced the larger DA1 response to ipsi-first motion observed in vivo (Figure 5t).

Thus, measured DA1 temporal filters plus a simple output nonlinearity are sufficient to generate ob-served ALPN direction selectivity, while leaving open contributions from the broader recurrent circuitry of the antennal lobe (Salman et al., 2026). In combination with the pharmacology, silencing data, and con-nectome analyses, this supports a model for the circuit in which the neuron il3LN6 supplies delayed and direction-selective inhibition to generate direction selectivity in the DA1 ALPN responses.

## DISCUSSION

Bilateral olfaction provides spatial information through two distinct but complementary comparisons: gradients report which side receives higher intensity odor input, whereas odor motion reports the direction of movement of odor signals across the antennae. Both cues guide source localization, homing, and plume tracking across animals, from mammals and sharks to migratory fish, lobsters, and insects (Borst and Heisenberg, 1982; Catania, 2013; Esquivelzeta Rabell et al., 2017; Gardiner and Atema, 2010; Kadakia et al., 2022; Kennedy and Marsh, 1974; Mafra-Neto and Cardé, 1994; Moore et al., 1991; Porter et al., 2007; Quinn and Dittman, 1990; Rajan et al., 2006; Taisz et al., 2023; Vickers and Baker, 1994). In walking Drosophila, odor motion improves navigation in odor plumes (Kadakia et al., 2022), and motion and gradient cues shape navigation differently in plumes with distinct spatiotemporal statistics (Brudner et al., 2025). What remained unknown was where and how odor motion is computed from bilateral odor inputs. Here, we described how the *Drosophila* antennal lobe transforms non-direction-selective olfactory input into direction-selective signals at the first synapse in the olfactory system. These results expand the range of olfactory computa-tions occurring in the antennal lobe. They also turn a behaviorally relevant cue into a circuit-level variable: a direction-selective signal that can be followed from the antennal lobe to the pathways that guide navigation in a turbulent chemical world.

How motion cues are converted into behavior remains unknown. It is possible that downstream circuits integrate gradient and motion cues through a common lateralized olfactory signal. Static gradients and moving odor filaments are physically distinct, yet both can create a lateralized imbalance between antennal lobe outputs across hemispheres: gradients by more strongly activating ALPNs ipsilateral to the higher-concentration antenna (Gaudry et al., 2013; Taisz et al., 2023), and motion by more strongly activating ALPNs on the ipsi-first side. In both cases, the side with more activity tends to be closer to the odor source (Taisz et al., 2023) or odor plume midline (Brudner et al., 2025; Kadakia et al., 2022). A premotor circuit that reads out this hemispheric imbalance could therefore drive turns without having to classify the origin of the imbalance (gradient, odor, or both). In that case, the same circuit could steer flies up an attractive gradient (Borst and Heisenberg, 1982; Taisz et al., 2023) and against the apparent motion of an attractive odor filament (Kadakia et al., 2022). Because ALPNs carry motion signals to the lateral horn and mushroom body, odor motion could support innate orientation (Dolan et al., 2019; Fişek and Wilson, 2014; Frechter et al., 2019), become a learned feature (Aso et al., 2023; Sandoz and Menzel, 2001; Strube-Bloss et al., 2016; Zimmerman et al., 2026), or be combined with heading and goal signals in the central complex to generate steering commands (Kathman et al., 2024; Lanz et al., 2026; Matheson et al., 2022), even as these cues are integrated with self-motion, wind, and plume intermittency (Budick and Dickinson, 2006; Demir et al., 2020; Duistermars et al., 2009). In this view, the antennal lobe maps intensity gradients and motion direction onto shared lateralized signals, while glomerulus-specific downstream processing could determine how pheromone, food, and aversive channels weight these cues for valence-appropriate behavior.

Other animals that use bilateral olfaction for navigation may also compute odor motion, but where this computation could occur depends on where bilateral odor signals converge. In *Drosophila*, most ORNs project bilaterally to homologous antennal lobe glomeruli, allowing ipsi- and contralateral odor signals to interact at the first synapse in the olfactory system, where odor motion direction first emerges. Rodents have a contrasting organization: each naris feeds an ipsilateral olfactory bulb before bilateral information is exchanged through interbulbar and anterior olfactory nucleus pathways (Esquivelzeta Rabell et al., 2017; Kikuta et al., 2010), suggesting potential loci for odor motion signals in mice. In insects whose receptor neurons project mainly ipsilaterally, including bees, moths, mosquitoes, and locusts, odor motion signals may arise later, where antennal lobe output, projection or centrifugal neurons, or premotor pathways bring bilateral olfactory inputs together (Anton and Homberg, 1999; Flanagan and Mercer, 1989; Homberg et al., 1988; Jiang et al., 2024; Namiki and Kanzaki, 2016; Namiki et al., 2014; Singh et al., 2023; Suzuki, 1975; Takasaki et al., 2012). Because odor motion can provide useful directional information for animals navigat-ing odor plumes (Brudner et al., 2025; Kadakia et al., 2022), these signals might generically arise at early stages in processing where bilateral olfactory inputs interact.

Olfactory motion detection parallels visual motion detection at both algorithmic and circuit levels. Al-gorithmically, both sensory systems are sensitive to pairwise correlations in their inputs (Chen et al., 2023; Livingstone and Conway, 2003; Salazar-Gatzimas et al., 2016), a fundamental signature of motion (Potters and Bialek, 1994). Moreover, ALPNs are sensitive to static gradients (Taisz et al., 2023) as well as motion, a property that mirrors motion detectors in the fly eye (Agrochao et al., 2020). These shared algorithmic structures raise the possibility that olfactory motion detectors are tuned to plume statistics (Brudner et al., 2025), just as visual motion detectors appear tuned to natural scene statistics (Clark and Fitzgerald, 2024; Clark et al., 2014). At the circuit level, many visual systems generate direction selectivity by comparing spatially separated inputs over time, and using delayed inhibition to suppress responses to motion in the null direction (Barlow and Levick, 1965; Gruntman et al., 2018). A second well-defined example comes from the mammalian retina, where starburst amacrine cells have direction-selective neurites that provide asymmetric inhibition to direction-selective ganglion cells (Briggman et al., 2011; Euler et al., 2002; Fried et al., 2002).

The il3LN6-DA1 circuit uses a related organization in olfaction: il3LN6 is a local amacrine-like inhibitory neuron that responds preferentially to motion in the null direction of the downstream DA1 ALPNs. Its direction selectivity and delay both appear capable of generating direction-selective ALPN output.

Thus, motion detection circuits across modalities and animal phyla share a circuit logic in which local inhibitory neurons with delays and direction-biased responses convert temporally structured inputs across spatially separated sensors into direction-selective output. This shared solution suggests that motion detection may occupy a narrow circuit design space, determined by constraints on development, biophysics, stimulus statistics, and the structure of the motion detection problem.

## ACKNOWLEDGMENTS

We thank John Carlson, James Jeanne, Kristyn Lizbinski, and members of the Emonet and Clark labs for helpful discussions on this project. We thank Kristin Scott and Tzumin Lee for donating fly stocks. Stocks obtained from the Bloomington *Drosophila* Stock Center (NIH P40OD018537) were used in this study. We thank Paul Forscher for providing us with linear motorized stages. We thank Joel Greenwood and the Kavli Neurotechnology Core at Yale for support in designing experimental apparatus. This study was funded by the NIH BRAIN Initiative (1RF1NS132840). GMS was partially supported by CAPES (Brazil) – Finance Code 001.

## AUTHOR CONTRIBUTIONS

GMS, HV, TE, and DAC conceived experiments and interpreted results. GMS, with input from all authors, de-signed and built the optogenetic and odor stimulus apparatus. GMS obtained responses to real odors, iden-tified glomeruli, performed behavioral experiments, performed connectomic analyses, and characterized the computational model. GMS and HV designed and obtained imaging pilot data using optogenetic moving bar, noise, unilateral, and naturalistic stimuli. GMS obtained publication data for moving bar, unilateral, and naturalistic stimuli. HV obtained publication data for drifting gratings, correlated noise, and noise stimuli.

GMS prepared figures for publication and wrote the first draft of the manuscript, which all authors edited.

## METHODS

### Experimental Design Overview

All experiments used adult mated female *Drosophila melanogaster*. Flies used for imaging and behavioral experiments were 2–6 and 3–6 days old, respectively. The biological replicate was the fly unless ROI, glomerulus, or trajectory is explicitly stated as the unit of analysis. ROI-level analyses were computed within flies for fly-level tests unless otherwise stated. Behavioral trajectory-level analyses used trajectories as the statistical unit. Sample sizes are reported as flies unless stated otherwise. Recordings were excluded before analysis if stimulus timing or imaging alignment failed, brain motion precluded ROI extraction, or inter-trial response correlation was below 0.25, unless otherwise noted. Stimuli and experimental conditions were randomized and interleaved within experiments where applicable. Analysis-specific details, including exclusion criteria and statistical units, are described in the relevant sections below.

### Fly strains and rearing

Flies were reared at 25°C and 50% relative humidity on a 12 h:12 h light-dark cycle. Flies were reared on standard agarose food (Archon D2 or Lab-Express 7033).

Experimental flies were sorted under *CO*_2_ anesthesia before experiments. For behavioral experiments, flies were sorted at least 72 h before testing. For imaging experiments, flies were 2–6 days post-eclosion, sorted at least 24 h before imaging, and immobilized on ice immediately before surgical preparation. For optogenetic experiments, flies were fed all-trans-retinal (ATR) at 1 mM in water, diluted from from a 100 mM ATR stock in DMSO and without food, for 24 h before experiments. After ATR was added, vials were covered with foil and light exposure was minimized.

### Fly stocks

Canton S and Orco-Gal4, gifts from John Carlson; GH146-Gal4 (Berdnik et al., 2008) (RRID: BDSC_91812); nSyb-Gal4 (Pauli et al., 2008) (RRID: BDSC_51635); R57C10.AD (Dionne et al., 2018) (RRID: BDSC_70746); VT046100.DBD (Tirian and Dickson, 2017) (RRID: BDSC_75076); UAS-GCaMP6f (Chen et al., 2013) (RRID: BDSC_42747); LexAop-CsChrimson.tdTomato (Klapoetke et al., 2014) (RRID: BDSC_82183); empty split-Gal4 control (Hampel et al., 2015) (RRID: BDSC_79603); Orco-LexA (Lai and Lee, 2006); and UAS-shibire^t^*^s^* (w; UAS-shibire^t^*^s^*; +) (Kitamoto, 2001).

### Genotypes by experiment

(Figure 1), pan-neuronal imaging: w / w; UAS-GCaMP6f / +; nSyb-Gal4 / +. Figure 2c-2g, pan-neuronal imaging: w / w; UAS-GCaMP6f/+; Orco-LexA, LexAop-CsChrimson / nSyb-Gal4. Figure 2h-2i, ALPN imaging: w / w; UAS-GCaMP6f / GH146-Gal4; Orco-LexA, LexAop-CsChrimson / +. Figure 3, ALPN imaging: w / w; UAS-GCaMP6f / GH146-Gal4; Orco-LexA, LexAop-CsChrimson / +. ORN imaging: w / w; UAS-GCaMP6f / Orco-Gal4; Orco-LexA, LexAop-CsChrimson / +. Figure 4, DA1 ALPN imag-ing: w / w; UAS-GCaMP6f / GH146-Gal4; Orco-LexA, LexAop-CsChrimson / +. Figure 5k-5n and 5q, il3LN6 imaging: w / w; UAS-GCaMP6f / R57C10.AD; Orco-LexA, LexAop-CsChrimson/VT046100.DBD. Figure 5g-5j and Extended Data Fig. 9-10, il3LN6 silencing behavior: w / w; UAS-shibire^t^*^s^* / R57C10.AD; Orco-LexA, LexAop-CsChrimson / VT046100.DBD. Control silencing behavior: w / w; UAS-shibire^t^*^s^* / p65.AD; Orco-LexA, LexAop-CsChrimson / GAL4.DBD.

### Two-photon imaging

For *in vivo* imaging, flies were food-deprived for 24-48 h in polystyrene vials (Genesee #32-110) capped with Flugs (Genesee #49-101), with water supplied from a soaked piece of Flugs at the bottom of the vial. For surgical preparation, flies were cold anesthetized and head-fixed in a metal shim with UV-curable glue (Liquid Plastic Adhesive, RapidFix UV). The brain was exposed by removing the cuticle above the anten-nae and between the eyes, followed by removal of visible fat tissue and trachea. To reduce brain motion, muscle M16 (Jayaraman and Laurent, 2007) was transected and the mouthparts and legs were fixed with UV-curable glue. The exposed brain was submerged in standard oxygenated sugar-saline imaging solution (Tanaka and Clark, 2022; Wilson et al., 2004).

Imaging was performed at room temperature 20 °C ± 1.5 °C on a two-photon microscope (Hyper-Scope, Scientifica) using a 20x water-immersion objective (N20X-PFH XLUMPNFL, 1.00 NA, 2 mm working distance, Olympus). GCaMP6f was excited at 930 nm with a femtosecond Ti:Sapphire laser (Mai Tai eHP, SpectraPhysics). Laser power at the sample was kept below 40 mW. Emitted light was filtered with two 515/30 filters in series (Semrock FF01-515/30-25). Image acquisition was controlled with ScanImage (Pologruto et al., 2003). All functional imaging used single imaging planes. Frame size and zoom were adjusted for each preparation and field of view; frame rates were 6 or 13.5 Hz depending on field of view.

Stimulus timing was aligned to the imaging data using a synchronization LED driven by the same command signal as the stimulus and recorded in the imaging data. The LED trace was extracted from each recording and used to identify stimulus onset times for trial segmentation and response alignment.

Ipsilateral and contralateral stimulus labels were defined relative to the hemisphere being imaged. Be-fore pooling across preparations, left and right antenna activation streams were mapped to ipsilateral and contralateral coordinates using the recorded imaging hemisphere. Each stimulus trial type was repeated five times unless stated otherwise.

### Odor and optogenetic stimulation

#### Moving odorized ribbon stimuli

Custom glass pipettes (146 mm length, 1.5 mm tip inner diameter) were fitted with miniature three-way solenoid valves (LHDA 1231515H, The Lee Company) for odor delivery. The pipette tip was positioned ≈ 1 cm from the fly. Valve inlets were placed 10 cm from the pipette tip.

Airflow was controlled with mass-flow controllers (Alicat Scientific). Total flow to the fly was 1 L min*^−^*^1^, with 50 mL min*^−^*^1^ odorized flow carried in clean air. Pipettes were mounted on motorized linear stages (TRB25CC, Newport; DDS100, Thorlabs). The odorant was undiluted apple cider vinegar (5% acidity, Great Value, Walmart). Stimuli lasted 20 s and were interleaved with 5 s intervals of clean air at 1 L min*^−^*^1^.

Each 20 s trial contained eight sequential odorized ribbon presentations, four ipsilateral-first and four contralateral-first, with directions alternating every ≈ 1.5 s as the stage moved back and forth at 10 mm s*^−^*^1^. Depending on the hemisphere being imaged, the first odorized ribbon presentation was ipsilateral-first or contralateral-first.

#### Optogenetic olfactory stimuli

Independent optogenetic activation of each antenna was achieved by positioning fiber-optic cannula tips (50 µm core, 0.22 NA, Doric Optical Probe Tip) as close as possi-ble, immediately adjacent to the antennae. CsChrimson was expressed in all olfactory receptor neurons (*Orco*-Gal4), and activated with 625 nm LEDs (M625F2, Thorlabs). Data acquisition hardware (T7, LabJack) controlled stimulus timing and LED activation, and photodiode recordings verified stimulus delivery. LED power at the left and right antennae was independently calibrated at the antennal plane to 3 µW mm*^−^*^2^ with a PM100D power meter (Thorlabs) and variable optical attenuator (VOAMMF, Thorlabs), and was held con-stant unless stated otherwise. The fiber optic tips were positioned with micromanipulators (Drummond Sci-entific).

For unilateral antenna activation experiments, each trial lasted 5 seconds, and each antenna re-ceived three 400 ms light pulses separated by 800 ms. For optogenetic moving bars, ipsilateral-first and contralateral-first stimuli were generated by sequentially activating ORNs in each antenna with a fixed inter-antenna delay. Each motion trial contained three optogenetic moving bars in a single direction. For each bar, each antenna received a 400 ms pulse, so total stimulation was matched across directions. Simulta-neous bilateral activation was used as a no-motion control where indicated. The standard inter-antenna delay was 200 ms unless a delay was experimentally varied. Imaging stimulus trials were separated by 5 s interleaves unless stated otherwise.

#### Delay tuning (**Figure 4a**)

Ipsilateral-first and contralateral-first moving-bar sequences were presented with inter-antenna delays of 25, 50, 100, 200 and 350 ms. Each bar stimulated each antenna for 400 ms, so total stimulation was matched across delays.

#### Direction-switching stimuli (**Figure 4j**)

Moving bars were presented sequentially, with motion direc-tion alternating between ipsilateral-first and contralateral-first on each successive presentation. The switch interval was defined as the time from the offset of the trailing bar to the onset of the next bar and was var-ied across 83, 250, 333 and 417 ms. For presentation *j*, we computed a signed DSI,

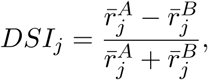

where *A* denotes the trace that began with ipsilateral-first motion and *B* denotes the trace that began with contralateral-first motion. Because the two traces exchanged motion direction after each switch, the sign of *DSI_j_* alternated across presentations. Direction-switching strength was computed for each fly as the mean absolute DSI, ⟨|*DSI_j_*|⟩, across presentations 2–4.

#### Correlated noise stimuli (**Figure 4e**)

Binary LED commands were sampled from a Bernoulli distri-bution with *P* (ON) = 0.5, updated at 50 Hz (20 ms frames), and presented in 3 s epochs separated by 3 s interleaves with no stimuli. ON frames drove the LED and OFF frames turned the LED off. Inter-antenna de-lays were imposed by shifting one antenna stream by 0, 1, 2, 4, 6 or 8 frames, corresponding to 0, 20, 40, 80, 120 or 160 ms delays. Positive-correlation stimuli used delayed copies of the same binary sequence at the two antennae, whereas negative-correlation stimuli used delayed inverted copies, with ON and OFF frames exchanged at one antenna. Ipsi-leading and contra-leading conditions were generated by swapping the antenna receiving the leading sequence. For each epoch, one sequence was randomly selected from 100 seeded binary random sequences.

#### Moving sinusoid stimuli (Extended Data Fig. 7)

Apparent motion was generated from two sinu-soidal signals, a reference signal *S*_ref_ = sin(2*πft*) and a lagged signal *S*_lag_ = sin(2*πft* − *ϕ*), where *f* is temporal frequency and *ϕ* is phase lag. The phase lag was set by *ϕ* = 2*π*Δ*_x_*/*λ*, where Δ*_x_* is the spatial sepa-ration between antennal receptive fields (≈ 250 µ*m*) and *λ* is spatial wavelength. Stimuli used spatial wave-lengths of 4Δ*_x_* and 16Δ*_x_*, corresponding to phase lags of *π*/2 and *π*/8, respectively, with temporal frequen-cies of 0.25, 0.5, 1, 2 and 4 Hz. Sinewave LED commands were offset and scaled as *I*(*t*) = *I*_max_(*S*(*t*) + 1)/2, so each antenna’s LED command ranged from a floor of 0 to *I*_max_. *I*_max_ was the calibrated antenna-plane ir-radiance of 3 µW mm*^−^*^2^, measured independently for the left and right LEDs. Motion direction was reversed by swapping which antenna received the phase-lagged signal.

#### Naturalistic stimuli (**Figure 4n**)

We used a naturalistic odor waveform from Kadakia et al. (Kadakia et al., 2022), originally measured using a photoionization-detector (PID) in earlier experiments (Gorur-Shandilya et al., 2017), as an intermittent odor stimulus. Analyses here used the first 35 s from the PID-recorded trace. The PID waveform was log-transformed, normalized, downsampled to 50 Hz, and used to drive optogenetic stimulation in the two antennae. The same waveform was presented to both antennae for the zero-shift condition or to one antenna with imposed temporal shifts of 1, 2, 4 and 8 frames at 50 Hz, corresponding to 20, 40, 80 and 160 ms. Ipsi-first and contra-first conditions were generated by swapping which antenna received the leading waveform.

#### Odor panel for glomerulus identification

For glomerulus identification, a custom glass pipette was fitted with six miniature three-way solenoid valves (LHDA 1231515H, The Lee Company). The valve closest to the tip was placed 7 cm from the pipette tip and subsequent valves were separated by 1 cm. Airflow was controlled as previously described. Six odorants were presented as a reference panel to label glomerular landmarks (Wang et al., 2003). Odorants were purchased from MilliporeSigma and diluted to 10*^−^*^3^ v/v in paraffin oil (76235, MilliporeSigma). Each odorant used dedicated PTFE-coated tubing (McMaster) and a dedicated valve to prevent cross-contamination. Odors were presented for 5 s with a 10 s inter-trial clean air interval. The odor panel contained pentyl acetate, butyl acetate, 3-hexanone, propyl acetate, methyl sal-icylate, and ethyl 3-hydroxybutyrate. Antennal lobe responses were imaged along the dorsal-ventral axis at up to seven planes: z ≈ 5, 15, 25, 40, 55, 80, and 105 µm, where z ≈ 0 was the dorsal-most surface.

### Behavioral assay

Behavioral experiments were performed in the walking arena described previously (Kadakia et al., 2022). The arena was 270 mm by 180 mm by 10 mm, with top and bottom glass surfaces and acrylic sidewalls. Flies were food-deprived for 72 h in polystyrene vials (Genesee #32-110) capped with Flugs (Genesee #49-101), with water supplied from a soaked piece of Flugs at the bottom of the vial. Fly vials were placed in a temperature-controlled chamber set to 30°C for 30 minutes before experiments. Approximately 30 flies were mouth aspirated into the behavioral arena and allowed to acclimate for 2 min before experiments.

The behavioral arena was kept at 29-30°C using temperature-controlled heating pads (Heat Pad 24W with thermostat, REPTI ZOO). Experiments were recorded at 60 Hz with an IR camera (GS3-U3-41C6NIR-C, FLIR), and tracked with custom Python scripts. Optogenetic stimuli were delivered with a DMD projector (LightCrafter 4500, Texas Instruments) mounted above the arena, using only the red channel (central wave-length 627 nm, 4.25 µW mm*^−^*^2^) at 60 Hz. Experiments were performed during the flies’ circadian active windows, within 3 h after lights-on or before lights-off.

#### Moving bars experiments (**Figure 5g**)

For il3LN6 silencing behavior, experimental and control flies were tested at 30°C to activate shibire^t^*^s^*, a temperature-sensitive dynamin allele that blocks synaptic vesicle recycling (Kitamoto, 2001). Control flies carried the empty split-Gal4 driver genotype, and experimental flies carried the il3LN6 split-Gal4 driver genotype. Flies were tested without wind. The stimulus consisted of 5 mm wide optogenetic bars moving at 15 mm s*^−^*^1^ (Kadakia et al., 2022). Bars were presented in alternating 5 s stimulus and 5 s blank blocks for 30 s total. The two motion directions were collected as separate +15 and −15 mm s*^−^*^1^ recordings and aligned during analysis.

#### Straight-ribbon experiments (Extended Data Fig. 10)

Flies were tested with constant laminar flow at 150 mm s*^−^*^1^ (Kadakia et al., 2022) and 30°C. Four stationary red-light ribbons were projected into the arena while flies walked freely. The ribbons were 5 mm wide in the cross-wind direction and spanned the arena length along the wind axis. Each recording lasted 60 s. Control flies carried the empty split-Gal4 driver genotype, and experimental flies carried the il3LN6 split-Gal4 driver genotype.

### Behavioral analysis

Tracked trajectories were segmented by track identity using centroid-based tracking (Kadakia et al., 2022) and filtered before analysis. Track identities could terminate and restart, so included trajectories were used as the statistical unit; multiple trajectories from the same fly could contribute independently, and no session-level averaging was applied. Moving-bar analyses included only trajectories with walking speeds of 2–25 mm s*^−^*^1^ that lasted at least 10 s.

For each bar motion direction, fly headings were rotated into a stimulus-centered reference frame in which 0° indicated orientation against bar motion and 180° indicated orientation with bar motion. Rotated headings were wrapped to [−180°, 180°) and folded by absolute value onto the range 0–180°. The two bar motion directions were then pooled, and folded headings were summarized in 24 bins from 0 to 180°.

#### KL-divergence analysis (Extended Data Fig. 9

Headings were folded from 0–180° and divided into 24 equal 7.5° bins. For each trajectory, we calculated the fraction of heading samples in each angular bin. These fractions were averaged across trajectories, giving every trajectory equal weight. For each genotype, we averaged the fraction of time spent in each angular bin across trajectories. Pairwise differences were quantified as the symmetric KL divergence, <u>^1^</u> [*D*_KL_(*P* ∥ *Q*) + *D*_KL_(*Q*∥ *P* )]. KL divergence was calculated using base-2 logarithms and reported in bits. A negligible 10*^−^*^12^ value was added to each bin before normalization to avoid undefined values for empty bins. Significance was assessed by randomly permuting genotype labels among trajectories 100,000 times while preserving group sizes. Permutation *P* values were calcu-lated as (*k* + 1)/(*N* + 1), where *k* is the number of shuffled KL values equal to or greater than the measured value and *N* = 100,000. This correction prevents *P* values of zero when no shuffle is as extreme as the mea-sured data.

For straight-ribbon analyses, trajectories required the same walking speed criterion and at least one encounter with an optogenetic ribbon. Marginal distributions show the y-positions of all analyzed trajecto-ries, binned into 140 bins, smoothed with a Gaussian kernel with *σ* = 2.5 bins, and normalized to the maxi-mum density within each panel.

### Image processing and ROI extraction

Imaging data were motion-corrected using custom rigid x-y frame-registration code and converted to fluo-rescence time series for each region of interest (ROI). For grid-based ROI extraction, ROIs were sampled on a 5 µm grid. For glomerular and local neuron analyses, ROIs were identified with response-driven watershed segmentation. For each ROI, local background fluorescence was estimated from unlabeled dim pixels in the same horizontal y-stripes as the ROI and subtracted before *dF* /*F*_0_ calculation. For each recording, mean interleave fluorescence was fit with *A* exp(−*t*/*τ* ) to estimate a shared photobleaching time constant. ROI-specific amplitudes were then estimated from interleave fluorescence after dividing out this exponential decay, yielding a time-varying baseline *F*_0_(*t*). Responses were expressed as fractional fluorescence change, *dF* /*F*_0_ = (*F* − *F*_0_(*t*))/*F*_0_(*t*), and are denoted as dF/F in figure axes. Baseline fluorescence for each stimulus was estimated from the final 1 s of the preceding interleave (no stimulus) epoch.

For response-driven ROI extraction, each pixel was assigned a response score from three terms: nor-malized mean response amplitude, local response similarity, and trial-repeat reproducibility. Normalized mean response amplitude was the mean stimulus-evoked response across trials for that pixel, normalized by its peak response. Local response similarity was the mean positive cosine similarity between a pixel’s trial-response vector and the trial-response vectors of its eight neighboring pixels. Trial-repeat reproducibil-ity was computed by splitting repeated presentations of each stimulus condition into two sets, averaging re-sponses within each set, and computing the cosine similarity between the two resulting stimulus-response sets for each pixel. The final score image was the product of normalized mean response amplitude, local re-sponse similarity, and trial-repeat reproducibility. This score image was segmented with an 8-connected watershed algorithm. Contiguous local maxima in the score image defined candidate ROI centers, and neighboring pixels were assigned to the corresponding candidate ROI. Candidate ROIs were restricted to high-scoring pixels by retaining pixels at or above the 75th percentile of scores across the image. ROIs with fewer than five pixels were discarded. DA1 analyses used a manually selected ROI identified from anatomy. Unless noted, the response metric was the mean baseline-subtracted *dF* /*F*_0_ over the stimulus epoch, and no temporal smoothing was applied.

ROIs were included in direction-selectivity analyses when they passed the inter-trial response reliability threshold. Reliability was computed as trial-to-trial response correlation across repeats of the same stimulus and across stimulus types. Threshold was *r* ≥ 0.35 for (Figure 1) odorized ribbon ROI inclusion and for glomerular ALPN and ORN-terminal analyses in (Figure 3). Threshold was *r* ≥ 0.25 for (Figure 2) unilateral antenna activation and optogenetic motion, for (Figure 4) except noise stimuli, and for il3LN6 imaging in Figure 5k-5n and 5q.

For glomerular ALPN and ORN-terminal analyses in (Figure 3), initial ROIs were grouped across recordings using response correlation, DSI, and spatial location, then manually adjusted, verified, and labeled by glomerulus. Glomerular identities were assigned from odor-panel landmarks, an antennal lobe atlas (Bates et al., 2020), and manual verification after aligning odor-response landmarks with atlas position and morphology. (Figure 3) response traces were smoothed with a Savitzky–Golay filter using a seven-sample window and a second-order polynomial. Raw traces were expressed as *dF* /*F*_0_. Normalized traces were scaled by the mean peak response for each glomerular label. DSI was computed for each ROI first, and ROI-level DSI values were then averaged across recordings within each glomerular label.

### Direction selectivity and response metrics

Direction selectivity was quantified with a single direction selectivity index (DSI) convention throughout the paper:

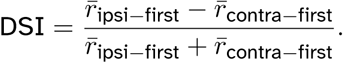

Here, *r̄*_ipsi*−*first_ and *r̄*_contra*−*first_ are mean responses for ipsilateral-first and contralateral-first motion stimuli. Ipsilateral-first and contralateral-first directions were defined relative to the imaged hemisphere. For sequential stimuli, the mean response was computed over the whole stimulus epoch. Positive DSI values indicate larger responses to ipsilateral-first motion.

For odorized ribbon analyses in (Figure 1), direction-selective ROIs were identified from responses to ipsilateral-first and contralateral-first odorized ribbon presentations. Each trial contained eight sequential response peaks with alternating motion directions. The first and last peaks were excluded because they had only one neighboring peak. For each of the six internal peaks, the response was compared with the mean of the two adjacent peaks, which both represented the opposite motion direction (Figure 1). To preserve the global DSI convention, each peak-wise DSI was direction-coded:

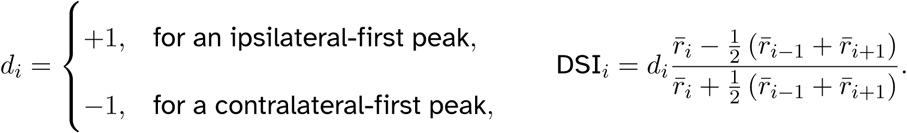

The ROI-level DSI was the mean of the six direction-coded peak-wise DSI values. Positive values therefore indicate larger responses to ipsilateral-first motion.

For shuffle-threshold analyses, the eight peak responses were randomly permuted 1,000 times within each ROI, and DSI was recomputed for each shuffle. The experiment-specific standard deviation of the shuffled DSI distribution (*σ*_shuffle_) was used as a threshold. For (Figure 1j) and (Figure 2g-2i), ROIs with |DSI| *> σ*_shuffle_ was computed for real and shuffled data in each fly.

For glomerular analyses in (Figure 3), DSI was first computed for each ROI and then averaged within fly and glomerular label, yielding one value per fly and glomerulus for statistical analysis.

For DA1 delay tuning in (Figure 4), DSI was computed at each delay with the global DSI convention from mean responses over the stimulus window. For correlated-noise stimuli, responses were averaged over 0–3 s after stimulus onset. Response differences were plotted as a function of imposed bilateral delay around zero offset. For per-delay summaries, responses were normalized within fly by the zero-delay response. Reverse-phi summaries compared the negative-correlation response difference with the positive-correlation response difference over matched delay ranges. For sinewave stimuli, responses were averaged over the stimulus window and summarized as ipsilateral-leading minus contralateral-leading responses across frequency and wavelength.

For naturalistic stimuli, whiffs were detected as peaks in the aligned stimulus waveform with a prominence threshold of 0.5 times the stimulus standard deviation. Peaks within 200 ms were merged by retaining the largest peak. Whiff responses were averaged from 100 ms before to 100 ms after each detected peak of calcium activity. Trial-level analyses subtracted the mean fluorescence during the first 3 s of each trial before whiff extraction. For each fly and imposed temporal shift, whiff response differences were averaged across detected whiffs as ipsilateral-leading minus contralateral-leading response. Delay-level means and bootstrap confidence intervals were computed across flies. The linear fit in (Figure 4q) was a descriptive least-squares line fit to delay-level mean whiff response differences across the four nonzero shifts (*r*^2^ = 0.856).

### Pharmacology

To block GABAergic inhibition in the antennal lobe, the GABA_A_ antagonist picrotoxin (#1128, Tocris Bio-science) and the GABA_B_ antagonist CGP 54626 hydrochloride (#1088, Tocris Bioscience) were applied in imaging saline. Picrotoxin stock was prepared in DMSO at 100 mM and stored at −20°C. CGP 54626 hy-drochloride was prepared fresh in ethanol on the day of the experiment. Before each experiment, antago-nists were diluted in fresh imaging solution to final concentrations of 20 µM picrotoxin and 50 µM CGP. The bath was replaced with either fresh imaging solution for saline controls or drug-containing imaging solution for GABA blockade. Responses were recorded 30 min after solution replacement.

Pharmacology experiments for DA1 ALPNs used the same genotype used for (Figure 4). Direction selectivity and response differences were computed as described above. For (Figure 5b-5d), responses were normalized by the mean peak response across trials, and the response difference was computed as ipsilateral-first minus contralateral-first response averaged over the stimulus window. Saline-control flies (n = 10) and GABA-antagonist-treated flies (n = 12) were compared with a two-sided Mann–Whitney U test.

### Linear response filters

Linear response filters were estimated using white-noise analysis (Chichilnisky, 2001; Matulis et al., 2020) using the timing-based kernel-extraction approach that permitted higher temporal resolution in the filters than the recording frame rate (Mano et al., 2019). Stimuli were updated at 50 Hz. On each frame, one of four activation patterns was presented: ipsilateral antenna only, contralateral antenna only, both antennae simultaneously, or blank. The stimulus contained 10 interleaved pairs of random-noise and fixed-noise blocks. Each random-noise block lasted 30 s and contained newly generated random activation patterns. Each fixed-noise block lasted 10 s and repeated the same random seed across repeats to estimate response reliability. The full stimulus lasted 400 s.

Filters were fit from *dF* /*F*_0_ traces and photodiode-aligned stimulus times. Left and right antenna acti-vation streams were converted to signed binary values and mapped to ipsilateral and contralateral streams using the recorded imaging hemisphere. For the ALPN filters used in the encoding model, the design ma-trix contained the current stimulus sample and 100 preceding stimulus samples, corresponding to a 2 s stimulus-history window at 50 Hz; no future stimulus samples were included. For each fluorescence sam-ple, the nearest stimulus sample was identified from the fitted stimulus clock (Mano et al., 2019); samples without a complete stimulus-history window were discarded, and the response vector was mean-centered before fitting. Ipsi- and contralateral filters were fit independently by ordinary least-squares regression. No regularization, deconvolution, post hoc filter normalization, scale factor, or offset was applied.

### Encoding model

A data-driven encoding model was built from DA1 ALPN response filters estimated by least squares fitting. Each recording produced one ipsilateral filter and one contralateral filter. The ALPN model filters were the mean ipsilateral and contralateral DA1 filters estimated across flies.

Moving-bar stimuli were represented as analytic binary traces at two point sensors corresponding to the ipsilateral and contralateral antennae. The two sensor positions were separated by 250 µm, approx-imating the distance between antennae in the fly. Bar speed was chosen to produce an ipsi-contra delay of 200 ms, matching *in vivo* recordings. Bar width was set so that each bar activated a sensor for 400 ms; each stimulus contained three sequential bars moving in the same direction, mirroring the experiments us-ing optogenetic moving bars in (Figure 2f). Measured filters were linearly interpolated from their native time base to a simulation time step 500-fold smaller, with the discrete filter sum preserved after interpolation.

Filters were not normalized. For each direction, ipsilateral and contralateral stimulus traces were convolved with the corresponding ALPN filters and summed to generate a linear response. In discrete time,

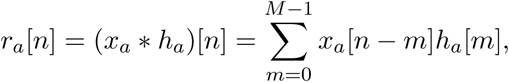

where *a* ∈ {ipsi, contra}, *x_a_* is the binary stimulus trace, and *h_a_* is the corresponding ALPN filter. Samples outside the stimulus trace were treated as zero, and the full convolution was truncated to the simulated time base. The summed linear drive was

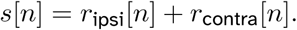

Model output was computed with a squared rectifying nonlinearity,

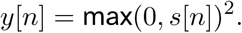

No scale factor or offset was fit. Model responses were summarized as the mean response over the stimulus span, and ipsilateral-first and contralateral-first responses were compared with the same DSI equation used for experimental responses.

### Connectome analysis

Connectome analyses used the FlyWire dataset (Dorkenwald et al., 2024; Schlegel et al., 2024). FlyWire queries used the FAFB public dataset with materialization 783 and FlyWire annotation tables. FlyWire source tables were generated in Python using CAVEclient queries to the flywire_fafb_public datastack at materialization 783, using the synapses_nt_v1 and fly_synapses_neuropil_v6 tables together with FlyWire annotation tables cached through fafbseg; annotations marked outlier_bio or outlier_seg were excluded.

To compute connectivity patterns, we used synapse counts as a proxy for connection weight (Scheffer et al., 2020). For each local neuron and glomerulus, ORN input was separated into ipsilateral and contralat-eral ORN synaptic input. ORN input laterality was computed as

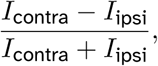

where *I*_ipsi_ and *I*_contra_ are ipsilateral and contralateral ORN input weights, respectively. Positive values there-fore indicate a contralateral ORN input bias. For each LN, a single ORN input preference was computed as an ORN-input-weighted mean across glomeruli, and averaged over LNs of the same type.

For scatter plots in (Figure 5e-5f), each circle represents one LN type. The shared y axis is the mean ORN input laterality for each LN type, computed as described above. Local neuron output to ALPNs was quantified per glomerulus as the summed LN-to-ALPN synaptic weight.

For the FlyWire bubble-matrix plot (Figure 8), bubble area represented the mean LN-to-ALPN output percentage across individual LNs of the same LN type and glomerulus. Total LN-to-ALPN synaptic weights were divided by the total output connections of that local neuron. Output fractions were multiplied by 100 to give the percentage of each local neuron’s output made onto ALPNs in that glomerulus.

Connectome summaries excluded non-olfactory VP glomeruli from antennal lobe glomerulus-level anal-yses. Individual LN and glomerulus-level analyses required at least 25 total ORN inputs per local neuron.

The bubble-matrix analysis further restricted LNs to those with ALPN output in at least three glomeruli be-fore applying the ORN-input filters. For each LN–glomerulus pair, the convergence score used in Figure 8 was defined as the contralateral ORN preference multiplied by the fraction of LN output onto ALPNs in that glomerulus and by log(1 + *W*_ALPN_), where *W*_ALPN_ is the LN-to-ALPN synapse count. Thus, high scores iden-tify LNs that receive strongly contralaterally biased ORN input and provide strong, concentrated output to ALPNs.

### Statistical analysis

Analyses were performed with custom Python scripts using NumPy (Harris et al., 2020), SciPy (Virtanen et al., 2020), Pandas (McKinney, 2010), and Matplotlib (Hunter, 2007). Statistical significance was defined as *P <* 0.05. Unless otherwise stated, paired tests were used for within-recording comparisons of matched ipsilateral/contralateral or ipsilateral-first/contralateral-first responses, and tests were two-sided. Exact paired sign tests were used for paired direction/count-style comparisons. Exact Wilcoxon signed-rank tests were used for paired il3LN6 response-amplitude comparisons in (Figure 5l) and (Figure 5n). Bootstrap con-fidence intervals were computed with 10,000 resamples with replacement.

Figure annotations use the following significance convention: ∗*P <* 0.05, ∗ ∗ *P <* 0.01, ∗ ∗ ∗*P <* 0.001, and ∗ ∗ ∗ ∗ *P <* 0.0001.

### Code and data availability

Figure source data, processed analysis tables, behavioral trajectory data, connectome-derived tables, model inputs and outputs, and analysis codes will be deposited in a public repository before publication. Public connectome data were obtained from FlyWire FAFB materialization 783.

---

**Extended Data Fig. 1:**
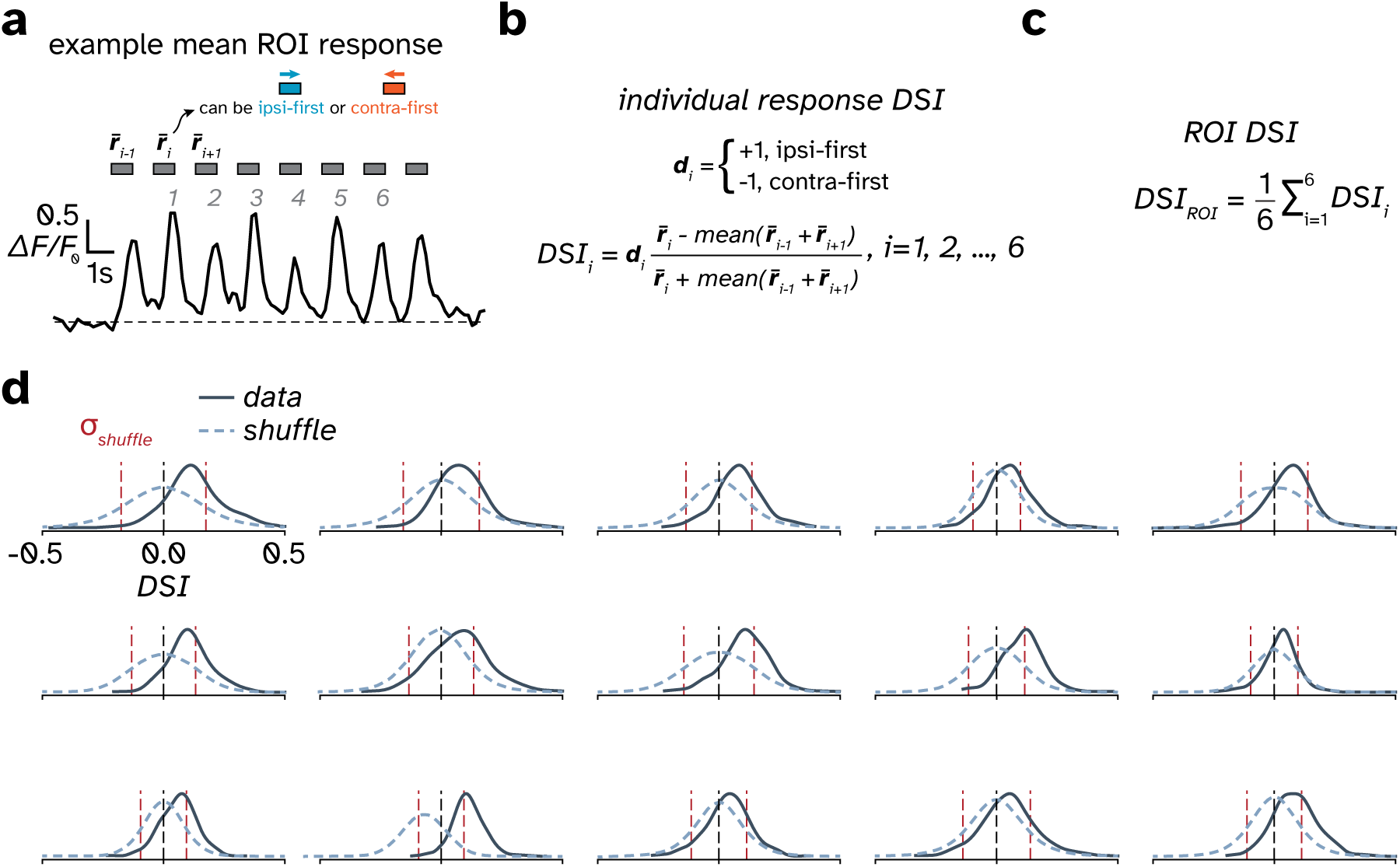
Quantification of lateral horn responses to moving odorized ribbons. **a.** Sample mean response from a lateral horn ROI to alternating ipsi-first and contra-first moving odorized ribbons, as described in Figure 1. Presentation direction was defined relative to the imaged hemisphere. See Direction selectivity and response metrics for details. **b.** Direction-coded DSI for each internal response peak. Each peak response was compared with the mean of its two neighboring peaks, which represented the opposite motion direction. The normalized contrast was multiplied by *d_i_* = +1 for ipsi-first peaks and *d_i_* = −1 for contra-first peaks, so positive values consistently indicated larger ipsi-first responses. **c.** ROI DSI, defined as the mean of the six direction-coded peak-wise DSI values. **d.** DSI distributions across lateral horn ROIs from pan-neuronal recordings and stimulus-label-shuffled controls. Each distribution represents one fly. Vertical dashed red lines mark ± 1 s.d. of the shuffled distribution.

**Extended Data Fig. 2:**
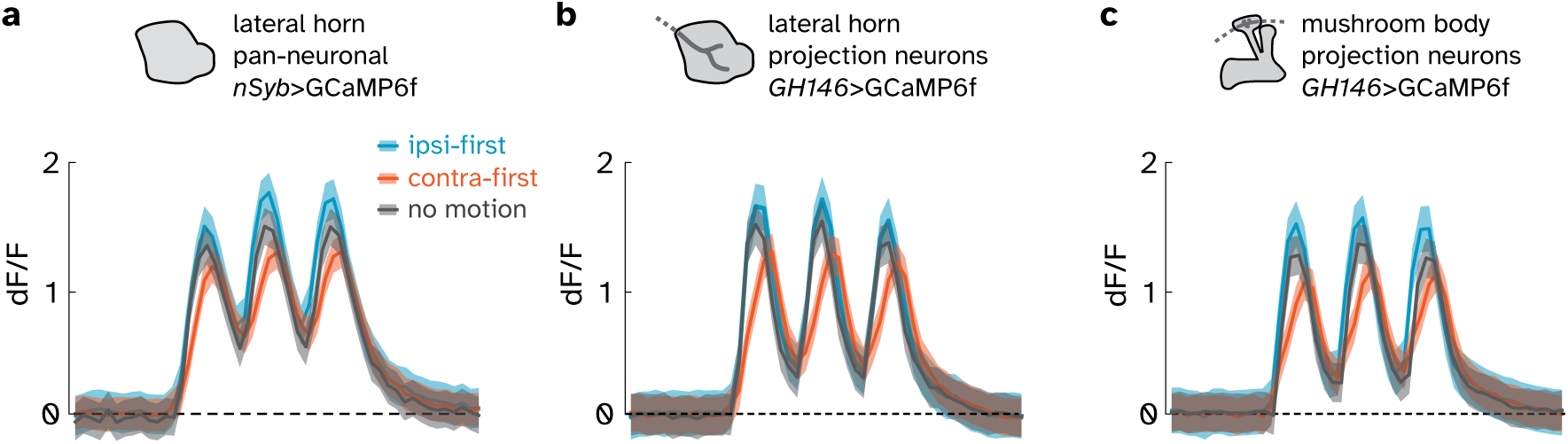
No-motion responses during optogenetic moving bar stimuli. Time-course responses (mean ± s.e.m.) to optogenetic moving bars, as in Figure 2g-2i, including the no-motion control condition. In the no-motion condition, the ipsilateral and contralateral antennae were activated simultaneously (Δ_delay_ = 0). See Odor and optogenetic stimulation in Methods for details. **a.** Pan-neuronal lateral horn responses; n = 13 flies. **b.** Antennal lobe projection neuron axon-terminal responses in the lateral horn; n = 11 flies. **c.** Antennal lobe projection neuron axon-terminal responses in the mushroom body calyx; n = 8 flies.

**Extended Data Fig. 3:**
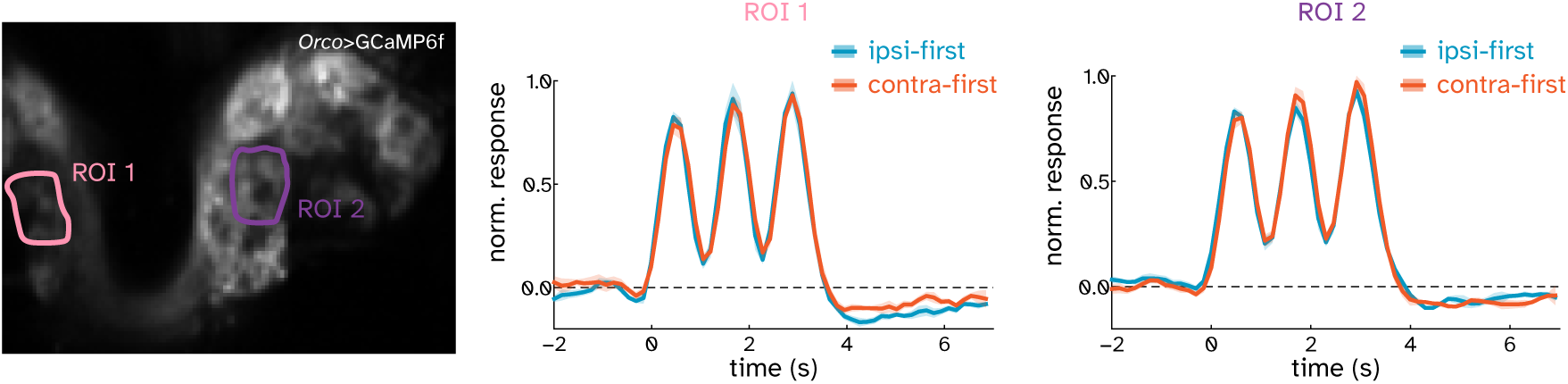
Matched ORN responses in opposing hemispheres to optogenetic moving bars. Normalized ORN terminal response time courses in *Orco*>GCaMP6f flies (see Methods for full genotype) during ipsi-first and contra-first moving bars, as in Figure 2f, measured from matched ROIs in the left (ROI 1) and right (ROI 2) hemispheres. For ROI 1, ipsi-first corresponds to left-to-right motion; for ROI 2, ipsi-first corresponds to right-to-left motion. Neither ROI showed a consistent preference for ipsi-first or contra-first motion. Traces show mean ± s.e.m. across five trials from one representative fly.

**Extended Data Fig. 4:**
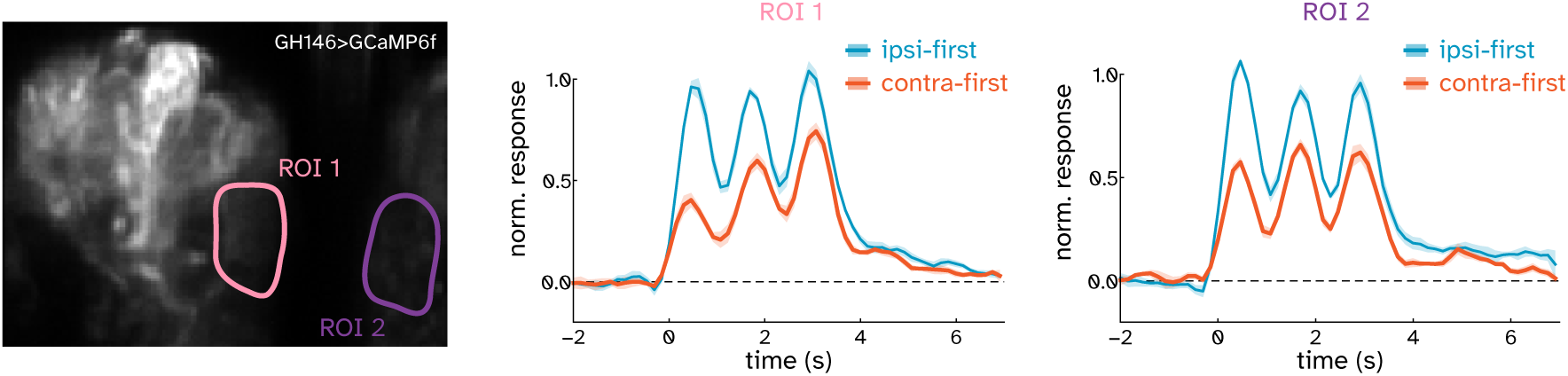
Matched ALPN responses in opposing hemispheres to optogenetic moving bars. Normalized ALPN dendrite response time courses in GH146>GCaMP6f flies (see Methods for full genotype) during ipsi-first and contra-first moving bars, as in Figure 2f, measured from matched ROIs in the left (ROI 1) and right (ROI 2) hemispheres. For ROI 1, ipsi-first corresponds to left-to-right motion; for ROI 2, ipsi-first corresponds to right-to-left motion. In both ROIs, ipsi-first motion evoked the larger response. Traces show mean ± s.e.m. across five trials from one representative fly.

**Extended Data Fig. 5:**
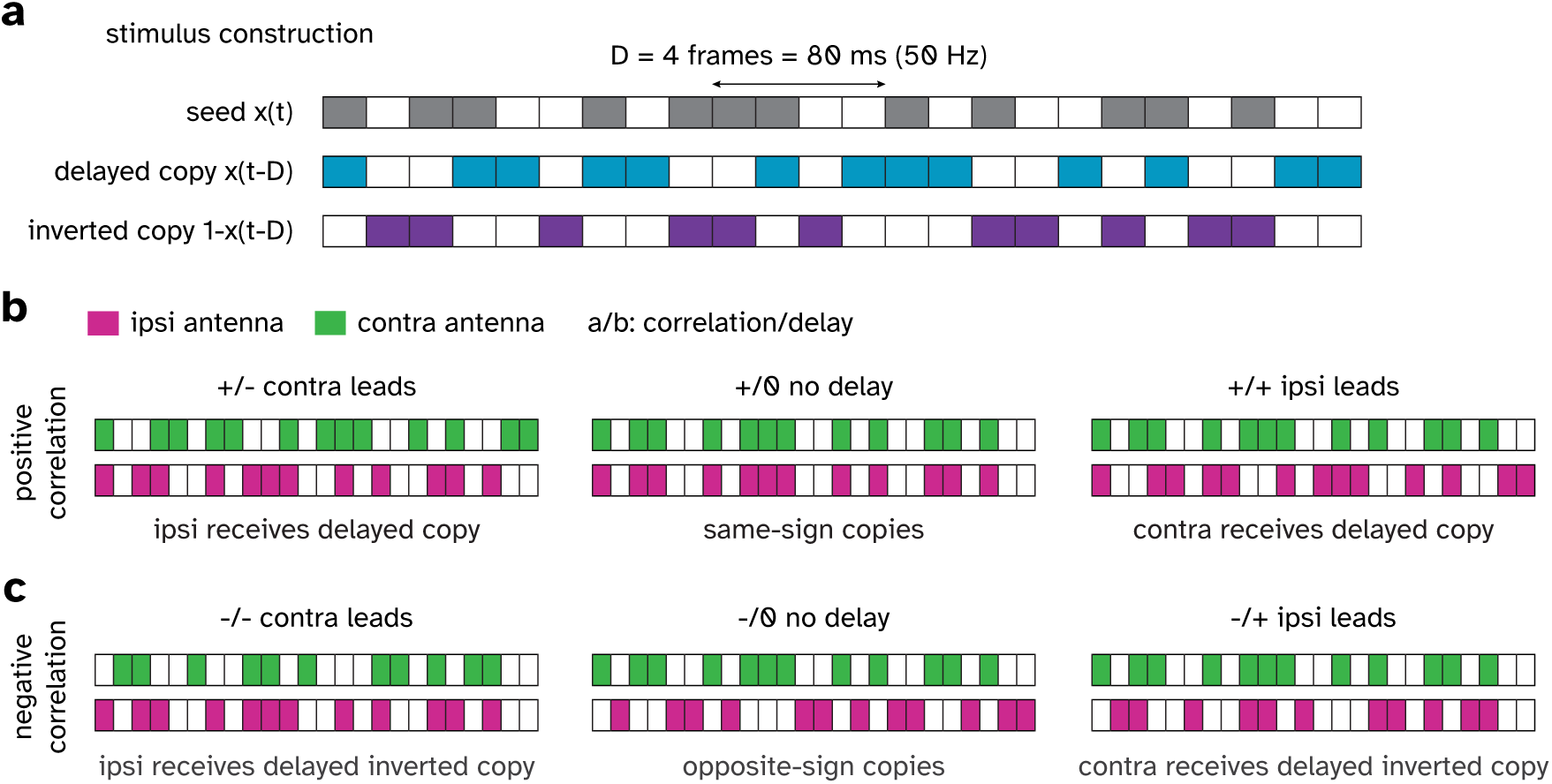
Correlated-noise stimulus construction. **a.** Binary optogenetic stimuli were generated from a random sequence *x*(*t*) updated at 50 Hz. Inter-antenna delay *D* was imposed by presenting a delayed copy, *x*(*t* − *D*), to one antenna. An inverted copy, 1 − *x*(*t* − *D*), was used for negative-correlation stimuli; the example shows *D* = 4 frames, corresponding to 80 ms (see Methods). Each rectangle represents one frame. White rectangles indicate no signal; colored rectangles indicate antenna activation. **b.** Sample time-series signal traces used to activate the ipsilateral and contralateral antennae. Positive-correlation stimuli used same-sign copies of the binary sequence at the two antennae. Each column shows one condition. The contra-leading and ipsi-leading conditions in this example used *D* = 1. The no delay condition used *D* = 0. Each rectangle represents one frame. White rectangles represent no signal; magenta rectangles represent ipsilateral antenna activation; green rectangles represent contralateral antenna activation; *a*/*b* nomenclature shows the imposed correlation and delay sign; e.g., +/− represents positive correlation and negative imposed delay. **c.** Negative-correlation stimuli used an inverted copy at one antenna. Signal traces as in **b**.

**Extended Data Fig. 6:**
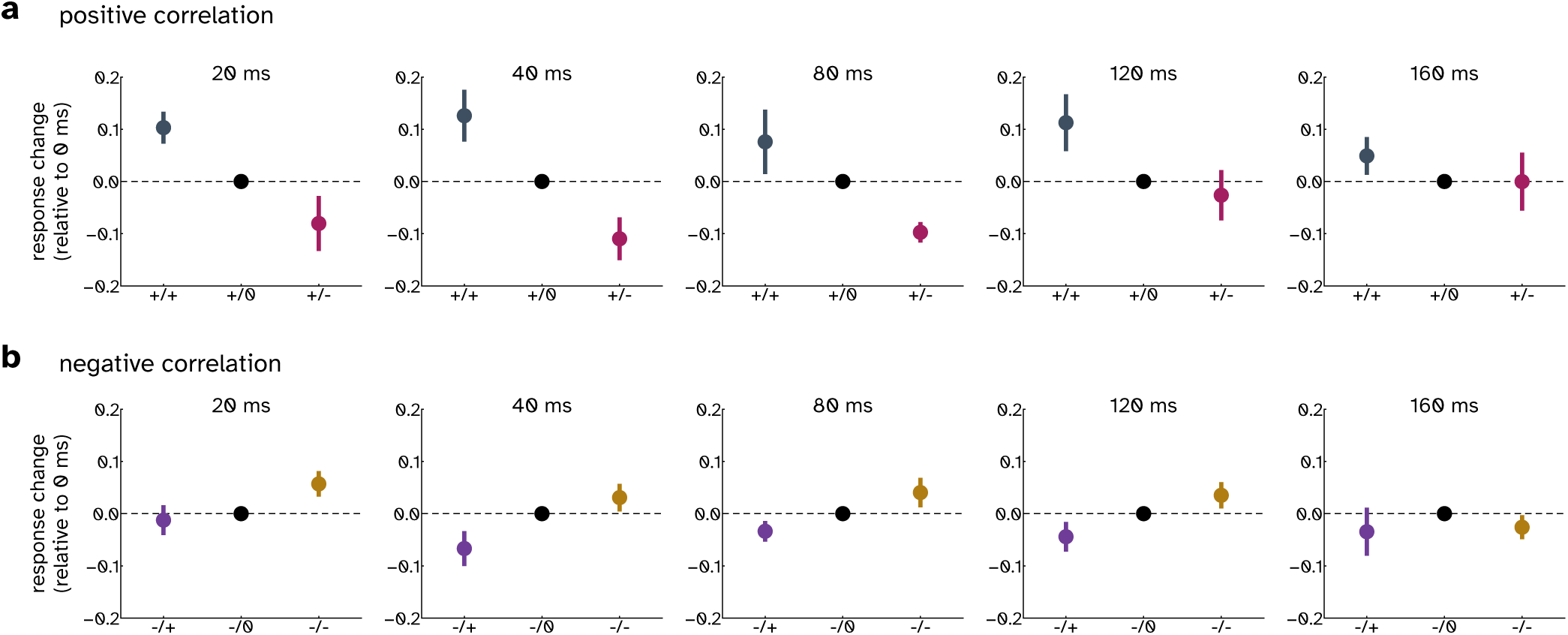
DA1 ALPN responses to correlated-noise stimuli across inter-antenna delays. **a.** Per-delay DA1 ALPN responses (mean ± s.e.m.) to positive-correlation stimuli. For each fly and delay, responses were normalized by the zero-delay response and plotted for ipsi-leading (+/+), zero-delay (+/0), and contra-leading (+/−) conditions (see Extended Data Fig. 5 for stimulus construction). Ipsi-leading and contra-leading responses were compared with exact two-sided paired sign tests at each delay. *P* values for 20, 40, 80, 120 and 160 ms were 0.0390625, 0.00390625, 0.00390625, 0.0390625 and 0.1796875, respectively; n = 9 flies. **b.** As in **a**, but for negative-correlation stimuli, shown as ipsi-leading (−/+), zero-delay (−/0), and contra-leading (−/−) conditions. Ipsi-leading versus contra-leading *P* values were 0.0703125, 0.0078125, 0.0703125, 0.2890625 and 0.2890625, respectively; n = 8 flies.

**Extended Data Fig. 7:**
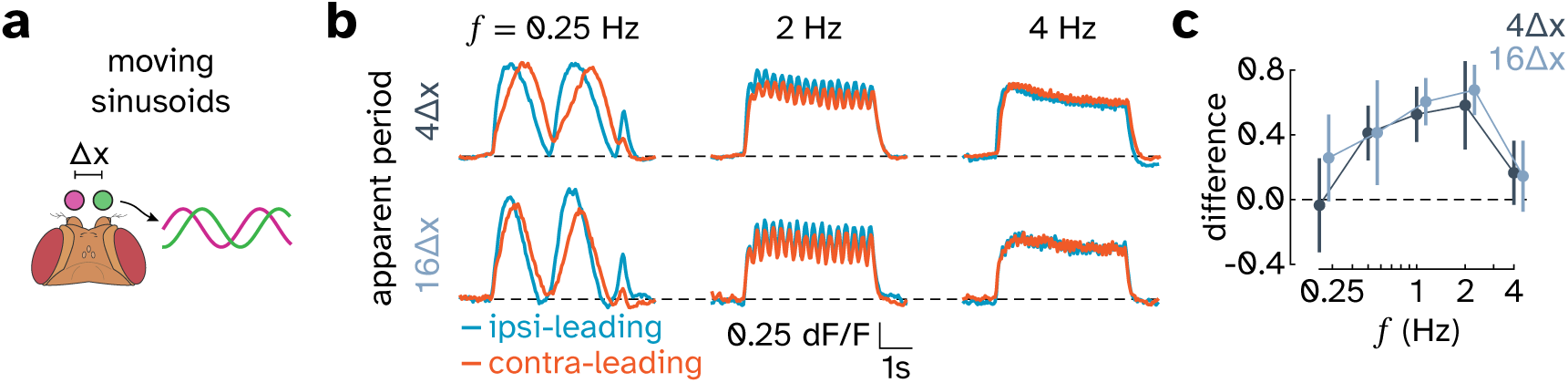
DA1 ALPN responses to moving sinusoids. Motion sensors can be broadly categorized as being tuned to the temporal frequency (TF) or the velocity of a stimulus. If a motion sensor is largely driven by pairwise intensity correlations and has simple filtering properties, then it will be sensitive to the TF when presented with moving sinusoids. The TF is defined as the ratio of the stimulus velocity to its spatial wavelength. We found that DA1 ALPN direction selectivity was strongest at TFs around 1-2 Hz, and weaker at low TFs. The responses were similar for both wavelengths tested, indicating that DA1 ALPNs were sensitive to stimulus TF and not to its velocity, similar to many visual motion detectors (Creamer et al., 2018; Foster et al., 1985; He and Levick, 2000). **a.** Moving sinusoid stimulus schematic. Bilateral sinusoidal stimulation varied temporal frequency *f* and spatial wavelength, expressed in antennal spacings (Δ*x*). **b.** DA1 ALPN responses to ipsi-leading and contra-leading moving sinusoids at selected temporal frequencies and spatial wavelengths of 4Δ*x* and 16Δ*x*. Traces show mean ± s.e.m. across flies; n = 7 flies for 4Δ*x*; n = 6 flies for 16Δ*x*. **c.** Time-averaged ipsi-leading minus contra-leading response difference as a function of temporal frequency for both spatial wavelengths. Circles show mean ± s.e.m. across flies; n = 7 flies for 4Δ*x*; n = 6 flies for 16Δ*x*.

**Extended Data Fig. 8:**
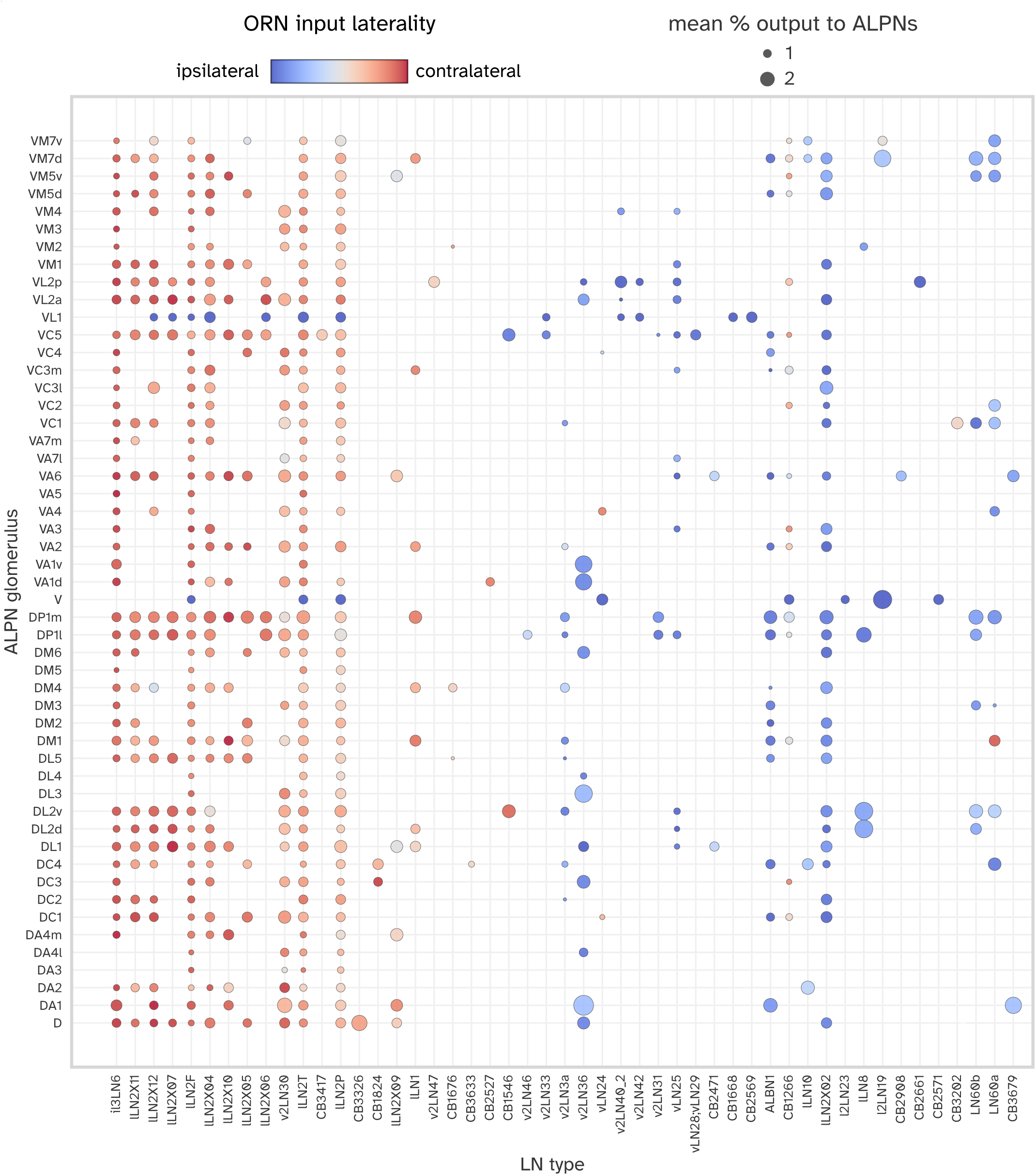
Connectomic input-output mapping of antennal lobe local neurons across glomeruli. FlyWire connectome analysis of antennal lobe local neuron (LN) types. Rows indicate ALPN glomeruli and columns indicate LN types, ordered by a convergence score that increases with contralateral ORN bias, the fraction of LN output directed to ALPNs, and LN-to-ALPN synapse count (see Methods). Each circle represents LN output from one LN type onto ALPNs in one glomerulus. Circle size indicates the mean percentage of that LN type’s output synapses onto ALPNs in the corresponding glomerulus. Circle color indicates the LN type’s mean ORN input laterality, computed as the normalized difference between contralateral and ipsilateral ORN input; blue indicates stronger ipsilateral ORN input and red indicates stronger contralateral ORN input.

**Extended Data Fig. 9:**
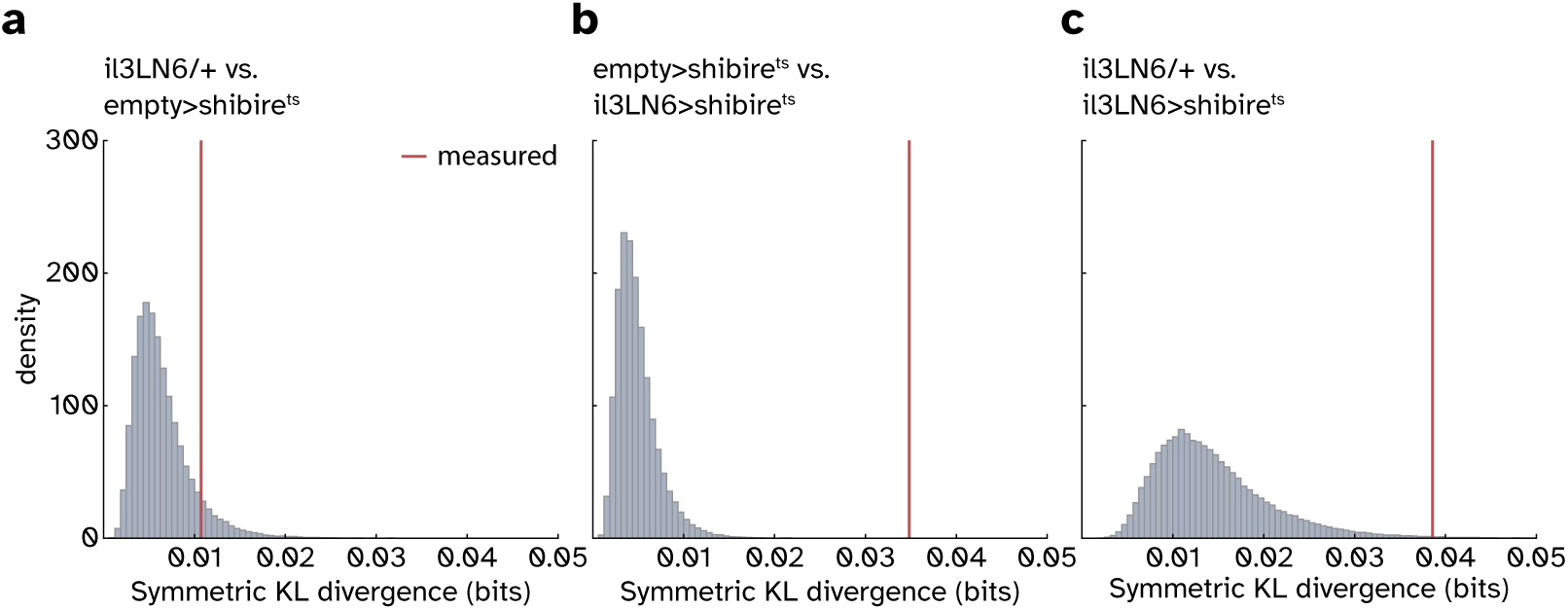
il3LN6 silencing reduces behavioral responses to optogenetic moving bars. Null distributions of pairwise symmetric KL divergence generated by 100,000 trajectory-level genotype-label permutations. Headings during the stimulus block were folded from 0–180° and divided into 24 bins. Vertical red lines indicate the measured KL divergence. *P* values are permutation tests. See Methods for details. **a.** il3LN6/+ versus empty>shibire^t^*^s^*; *P* = 0.0857. **b.** empty>shibire^t^*^s^* versus il3LN6>shibire^t^*^s^*; *P* < 0.0001. **c.** il3LN6/+ versus il3LN6/+; *P* = 0.00574.

**Extended Data Fig. 10:**
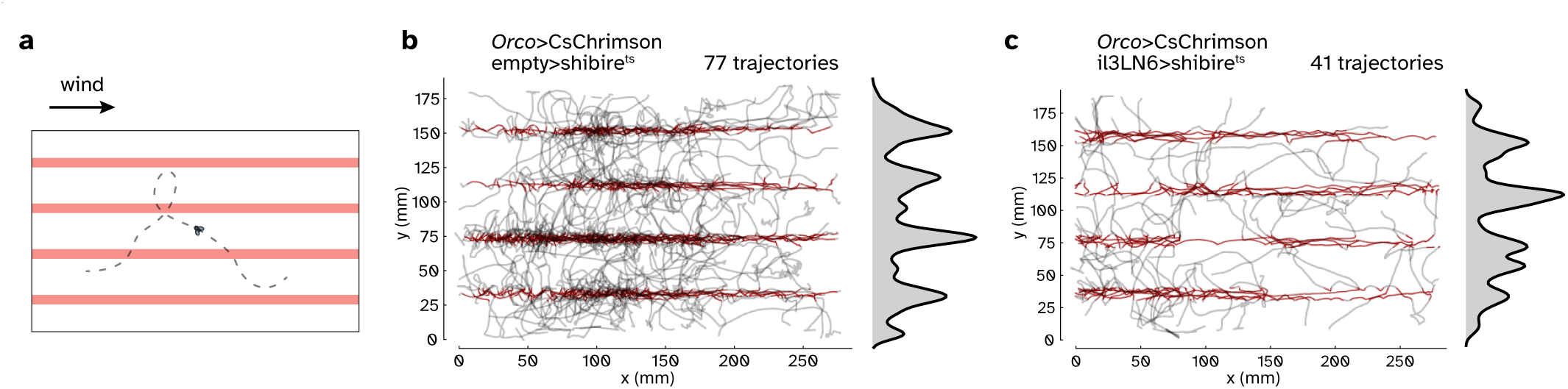
il3LN6-silenced flies track straight optogenetic odor ribbons in wind. **a.** Schematic of the straight-ribbon assay. Four stationary optogenetic odor ribbons were projected into the arena while wind flowed along the long axis of the ribbons. **b.** Trajectories of control empty>shibire^t^*^s^* flies during the straight-ribbon assay. Gray traces show retained walking trajectories; red trajectory segments mark periods where flies are within the ribbon area. Right marginals show the smoothed distribution of y positions across trajectory samples. Control flies and il3LN6-silenced flies both concentrated trajectories along the projected ribbons, indicating that il3LN6 silencing did not abolish optogenetic odor detection or odor-guided walking in wind. See Methods for trajectory inclusion criteria and marginal-density estimation. n = 77 trajectories. **c.** As in **b** but for il3LN6>shibire^t^*^s^* flies. n = 41 trajectories.

## REFERENCES

1. E. H. Adelson and J. R. Bergen. Spatiotemporal energy models for the perception of motion. JOSA A, 2(2):284–299, Feb. 1985. ISSN 1520-8532. doi: 10.1364/JOSAA.2.000284. URL https://opg.optica.org/josaa/abstract.cfm?uri=josaa-2-2-284.

2. M. Agrochao, R. Tanaka, E. Salazar-Gatzimas, and D. A. Clark. Mechanism for analogous illusory motion perception in flies and humans. Proceedings of the National Academy of Sciences, 117(37):23044–23053, Sept. 2020. doi: 10.1073/pnas.2002937117. URL https://www.pnas.org/doi/abs/10.1073/pnas.2002937117.

3. S. M. Anstis. Phi movement as a subtraction process. Vision Research, 10(12):1411–IN5, Dec. 1970. ISSN 0042-6989. doi: 10.1016/0042-6989(70)90092-1. URL https://www.sciencedirect.com/science/article/pii/0042698970900921.

4. S. Anton and U. Homberg. Antennal lobe structure. In B. S. Hansson, editor, Insect Olfaction, pages 97–124. Springer, Berlin, Heidelberg, 1999. doi: 10.1007/978-3-662-07911-9_5.

5. Y. Aso, D. Yamada, D. Bushey, K. L. Hibbard, M. Sammons, H. Otsuna, Y. Shuai, and T. Hige. Neural circuit mechanisms for transforming learned olfactory valences into wind-oriented movement. eLife, 12:e85756, Sept. 2023. ISSN 2050-084X. doi: 10.7554/eLife.85756. URL 10.7554/eLife.85756.

6. L. Badel, K. Ohta, Y. Tsuchimoto, and H. Kazama. Decoding of Context-Dependent Olfactory Behavior in Drosophila. Neuron, 91 (1):155–167, July 2016. ISSN 08966273. doi: 10.1016/j.neuron.2016.05.022. URL https://linkinghub.elsevier.com/ retrieve/pii/S089662731630201X.

7. H. B. Barlow and W. R. Levick. The mechanism of directionally selective units in rabbit’s retina. The Journal of Physiology, 178(3): 477–504, June 1965. ISSN 00223751. doi: 10.1113/jphysiol.1965.sp007638. URL https://onlinelibrary.wiley.com/doi/ 10.1113/jphysiol.1965.sp007638.

8. A. S. Bates, P. Schlegel, R. J. Roberts, N. Drummond, I. F. Tamimi, R. Turnbull, X. Zhao, E. C. Marin, P. D. Popovici, S. Dhawan, A. Jamasb, A. Javier, L. Serratosa Capdevila, F. Li, G. M. Rubin, S. Waddell, D. D. Bock, M. Costa, and G. S. Jefferis. Complete Connectomic Reconstruction of Olfactory Projection Neurons in the Fly Brain. Current Biology, 30(16):3183–3199.e6, Aug. 2020. ISSN 09609822. doi: 10.1016/j.cub.2020.06.042. URL https://linkinghub.elsevier.com/retrieve/pii/S0960982220308587.

9. R. Benton, J. Mermet, A. Jang, K. Endo, S. Cruchet, and K. Menuz. An integrated anatomical, functional and evolutionary view of the Drosophila olfactory system. EMBO Reports, 26(12):3204–3225, May 2025. ISSN 1469-3178. doi: 10.1038/s44319-025-00476-8. URL https://www.embopress.org/doi/full/10.1038/s44319-025-00476-8.

10. D. Berdnik, A. P. Fan, C. J. Potter, and L. Luo. MicroRNA processing pathway regulates olfactory neuron morphogenesis. Current biology: CB, 18(22):1754–1759, Nov. 2008. ISSN 0960-9822. doi: 10.1016/j.cub.2008.09.045.

11. A. Borst and M. Egelhaaf. Principles of visual motion detection. Trends in Neurosciences, 12(8):297–306, Jan. 1989. ISSN 01662236. doi: 10.1016/0166-2236(89)90010-6. URL https://linkinghub.elsevier.com/retrieve/pii/0166223689900106.

12. A. Borst and M. Heisenberg. Osmotropotaxis inDrosophila melanogaster. Journal of comparative physiology, 147(4):479–484, Dec. 1982. ISSN 1432-1351. doi: 10.1007/BF00612013. URL 10.1007/BF00612013.

13. K. L. Briggman, M. Helmstaedter, and W. Denk. Wiring specificity in the direction-selectivity circuit of the retina. Nature, 471(7337): 183–188, Mar. 2011. ISSN 1476-4687. doi: 10.1038/nature09818.

14. S. Brudner, B. Zhou, V. Jayaram, G. M. Santana, D. A. Clark, and T. Emonet. Fly navigational responses to odor motion and gradient cues are tuned to plume statistics, Apr. 2025. URL https://www.biorxiv.org/content/10.1101/2025.03.31.646361v1. Pages: 2025.03.31.646361 Section: New Results.

15. S. A. Budick and M. H. Dickinson. Free-flight responses of Drosophila melanogaster to attractive odors. Journal of Experimental Biology, 209(15):3001–3017, Aug. 2006. ISSN 0022-0949. doi: 10.1242/jeb.02305. URL 10.1242/jeb.02305.

16. K. C. Catania. Stereo and serial sniffing guide navigation to an odour source in a mammal. Nature Communications, 4(1):1441, Feb. 2013. ISSN 2041-1723. doi: 10.1038/ncomms2444. URL https://www.nature.com/articles/ncomms2444.

17. A. Celani, E. Villermaux, and M. Vergassola. Odor Landscapes in Turbulent Environments. Physical Review X, 4(4):041015, Oct. 2014. doi: 10.1103/PhysRevX.4.041015. URL https://link.aps.org/doi/10.1103/PhysRevX.4.041015.

18. J. Chen, C. M. Gish, J. W. Fransen, E. Salazar-Gatzimas, D. A. Clark, and B. G. Borghuis. Direct comparison reveals algorithmic similarities in fly and mouse visual motion detection. iScience, 26(10):107928, Oct. 2023. ISSN 2589-0042. doi: 10.1016/j.isci.2023.107928.

19. T.-W. Chen, T. J. Wardill, Y. Sun, S. R. Pulver, S. L. Renninger, A. Baohan, E. R. Schreiter, R. A. Kerr, M. B. Orger, V. Jayaraman, L. L. Looger, K. Svoboda, and D. S. Kim. Ultrasensitive fluorescent proteins for imaging neuronal activity. Nature, 499(7458): 295–300, July 2013. ISSN 0028-0836, 1476-4687. doi: 10.1038/nature12354. URL https://www.nature.com/articles/nature12354.

20. E. J. Chichilnisky. A simple white noise analysis of neuronal light responses. Network: Computation in Neural Systems, 12(2):199–213, 2001. doi: 10.1080/713663221.

21. Y.-H. Chou, M. L. Spletter, E. Yaksi, J. C. S. Leong, R. I. Wilson, and L. Luo. Diversity and wiring variability of olfactory local interneurons in the Drosophila antennal lobe. Nature Neuroscience, 13(4):439–449, Apr. 2010. ISSN 1097-6256, 1546-1726. doi: 10.1038/nn.2489. URL http://www.nature.com/articles/nn.2489.

22. D. A. Clark and J. E. Fitzgerald. Optimization in Visual Motion Estimation. Annual Review of Vision Science, Apr. 2024. doi: 10.1146/annurev-vision-101623-025432. URL https://www.annualreviews.org/content/journals/10.1146/annurev-vision-101623-025432.

23. D. A. Clark, J. E. Fitzgerald, J. M. Ales, D. M. Gohl, M. A. Silies, A. M. Norcia, and T. R. Clandinin. Flies and humans share a motion estimation strategy that exploits natural scene statistics. Nature Neuroscience, 17(2):296–303, Feb. 2014. ISSN 1546-1726. doi: 10.1038/nn.3600. URL https://www.nature.com/articles/nn.3600.

24. M. S. Creamer, O. Mano, and D. A. Clark. Visual Control of Walking Speed in Drosophila. Neuron, 100(6):1460–1473.e6, Dec. 2018. ISSN 1097-4199. doi: 10.1016/j.neuron.2018.10.028.

25. M. Demir, N. Kadakia, H. D. Anderson, D. A. Clark, and T. Emonet. Walking Drosophila navigate complex plumes using stochastic decisions biased by the timing of odor encounters. eLife, 9:e57524, Nov. 2020. ISSN 2050-084X. doi: 10.7554/eLife.57524. URL https://elifesciences.org/articles/57524. ZSCC: 0000012.

26. H. Dionne, K. L. Hibbard, A. Cavallaro, J.-C. Kao, and G. M. Rubin. Genetic Reagents for Making Split-GAL4 Lines in Drosophila. Genetics, 209(1):31–35, May 2018. ISSN 1943-2631. doi: 10.1534/genetics.118.300682.

27. M.-J. Dolan, S. Frechter, A. S. Bates, C. Dan, P. Huoviala, R. J. Roberts, P. Schlegel, S. Dhawan, R. Tabano, H. Dionne, C. Christoforou, K. Close, B. Sutcliffe, B. Giuliani, F. Li, M. Costa, G. Ihrke, G. W. Meissner, D. D. Bock, Y. Aso, G. M. Rubin, and G. S. Jefferis. Neurogenetic dissection of the Drosophila lateral horn reveals major outputs, diverse behavioural functions, and interactions with the mushroom body. eLife, 8:e43079, May 2019. ISSN 2050-084X. doi: 10.7554/eLife.43079. URL 10.7554/eLife.43079.

28. S. Dorkenwald, A. Matsliah, A. R. Sterling, P. Schlegel, S.-c. Yu, C. E. McKellar, A. Lin, M. Costa, K. Eichler, Y. Yin, W. Silversmith, C. Schneider-Mizell, C. S. Jordan, D. Brittain, A. Halageri, K. Kuehner, O. Ogedengbe, R. Morey, J. Gager, K. Kruk, E. Perlman, R. Yang, D. Deutsch, D. Bland, M. Sorek, R. Lu, T. Macrina, K. Lee, J. A. Bae, S. Mu, B. Nehoran, E. Mitchell, S. Popovych, J. Wu, Z. Jia, M. A. Castro, N. Kemnitz, D. Ih, A. S. Bates, N. Eckstein, J. Funke, F. Collman, D. D. Bock, G. S. X. E. Jefferis, H. S. Seung, and M. Murthy. Neuronal wiring diagram of an adult brain. Nature, 634(8032):124–138, Oct. 2024. ISSN 1476-4687. doi: 10.1038/s41586-024-07558-y. URL https://www.nature.com/articles/s41586-024-07558-y.

29. B. J. Duistermars, D. M. Chow, and M. A. Frye. Flies require bilateral sensory input to track odor gradients in flight. Current biology: CB, 19(15):1301–1307, Aug. 2009. ISSN 1879-0445. doi: 10.1016/j.cub.2009.06.022.

30. M. Egelhaaf, A. Borst, and W. Reichardt. Computational structure of a biological motion-detection system as revealed by local detector analysis in the fly’s nervous system. *Journal of the Optical Society of America. A*, Optics and Image Science, 6(7):1070–1087, July 1989. ISSN 0740-3232. doi: 10.1364/josaa.6.001070.

31. J. Esquivelzeta Rabell, K. Mutlu, J. Noutel, P. Martin Del Olmo, and S. Haesler. Spontaneous Rapid Odor Source Localization Behavior Requires Interhemispheric Communication. Current Biology, 27(10):1542–1548.e4, May 2017. ISSN 09609822. doi: 10.1016/j.cub.2017.04.027. URL https://linkinghub.elsevier.com/retrieve/pii/S0960982217304360.

32. T. Euler, P. B. Detwiler, and W. Denk. Directionally selective calcium signals in dendrites of starburst amacrine cells. Nature, 418 (6900):845–852, Aug. 2002. ISSN 0028-0836. doi: 10.1038/nature00931.

33. J. E. Fitzgerald, A. Y. Katsov, T. R. Clandinin, and M. J. Schnitzer. Symmetries in stimulus statistics shape the form of visual motion estimators. Proceedings of the National Academy of Sciences, 108(31):12909–12914, Aug. 2011. doi: 10.1073/pnas.1015680108. URL https://www.pnas.org/doi/full/10.1073/pnas.1015680108.

34. M. Fişek and R. I. Wilson. Stereotyped connectivity and computations in higher-order olfactory neurons. Nature Neuroscience, 17(2): 280–288, Feb. 2014. ISSN 1546-1726. doi: 10.1038/nn.3613. URL https://www.nature.com/articles/nn.3613.

35. D. Flanagan and A. R. Mercer. An atlas and 3-D reconstruction of the antennal lobes in the worker honey bee, Apis mellif-era l. (Hymenoptera: Apidae). International Journal of Insect Morphology and Embryology, 18(2–3):145–159, 1989. doi: 10.1016/0020-7322(89)90023-8.

36. K. H. Foster, J. P. Gaska, M. Nagler, and D. A. Pollen. Spatial and temporal frequency selectivity of neurones in visual cortical areas V1 and V2 of the macaque monkey. The Journal of Physiology, 365:331–363, Aug. 1985. doi: 10.1113/jphysiol.1985.sp015776. URL https://physoc.onlinelibrary.wiley.com/doi/10.1113/jphysiol.1985.sp015776.

37. S. Frechter, A. S. Bates, S. Tootoonian, M.-J. Dolan, J. Manton, A. R. Jamasb, J. Kohl, D. Bock, and G. Jefferis. Functional and anatomical specificity in a higher olfactory centre. eLife, 8:e44590, May 2019. ISSN 2050-084X. doi: 10.7554/eLife.44590. URL 10.7554/eLife.44590.

38. S. I. Fried, T. A. Münch, and F. S. Werblin. Mechanisms and circuitry underlying directional selectivity in the retina. Nature, 420 (6914):411–414, Nov. 2002. ISSN 0028-0836, 1476-4687. doi: 10.1038/nature01179. URL http://www.nature.com/ articles/nature01179.

39. J. M. Gardiner and J. Atema. The function of bilateral odor arrival time differences in olfactory orientation of sharks. Current biology: CB, 20(13):1187–1191, July 2010. ISSN 1879-0445. doi: 10.1016/j.cub.2010.04.053.

40. Q. Gaudry, E. J. Hong, J. Kain, B. L. de Bivort, and R. I. Wilson. Asymmetric neurotransmitter release enables rapid odour lateralization in Drosophila. Nature, 493(7432):424–428, Jan. 2013. ISSN 1476-4687. doi: 10.1038/nature11747. URL https://www.nature.com/articles/nature11747. Publisher: Nature Publishing Group.

41. S. Gorur-Shandilya, M. Demir, J. Long, D. A. Clark, and T. Emonet. Olfactory receptor neurons use gain control and complementary kinetics to encode intermittent odorant stimuli. eLife, 6:e27670, 2017. doi: 10.7554/eLife.27670. URL 10.7554/eLife.27670.

42. E. Gruntman, S. Romani, and M. B. Reiser. Simple integration of fast excitation and offset, delayed inhibition computes directional selectivity in Drosophila. Nature Neuroscience, 21(2):250–257, Feb. 2018. ISSN 1546-1726. doi: 10.1038/s41593-017-0046-4. URL https://www.nature.com/articles/s41593-017-0046-4. Publisher: Nature Publishing Group.

43. N. M. Grzywacz and F. R. Amthor. Robust directional computation in on-off directionally selective ganglion cells of rabbit retina. Visual Neuroscience, 24(4):647–661, July 2007. ISSN 1469-8714, 0952-5238. doi: 10.1017/S0952523807070666. URL https://www.cambridge.org/core/journals/visual-neuroscience/article/abs/robust-directional-computation-in-onoff-directionally-selective-ganglion-cells-of-rabbit-retina/5B8038FCBDAC96B1D5227F45696853C6.

44. J. Haag, W. Denk, and A. Borst. Fly motion vision is based on Reichardt detectors regardless of the signal-to-noise ratio. Proceedings of the National Academy of Sciences, 101(46):16333–16338, Nov. 2004. doi: 10.1073/pnas.0407368101. URL https://www.pnas.org/doi/full/10.1073/pnas.0407368101.

45. S. Hampel, R. Franconville, J. H. Simpson, and A. M. Seeds. A neural command circuit for grooming movement control. eLife, 4: e08758, Sept. 2015. ISSN 2050-084X. doi: 10.7554/eLife.08758. URL 10.7554/eLife.08758.

46. C. R. Harris, K. J. Millman, S. J. van der Walt, R. Gommers, P. Virtanen, D. Cournapeau, E. Wieser, J. Taylor, S. Berg, N. J. Smith, R. Kern, M. Picus, S. Hoyer, M. H. van Kerkwijk, M. Brett, A. Haldane, J. F. del Río, M. Wiebe, P. Peterson, P. Gérard-Marchant, K. Sheppard, T. Reddy, W. Weckesser, H. Abbasi, C. Gohlke, and T. E. Oliphant. Array programming with NumPy. Nature, 585 (7825):357–362, Sept. 2020. ISSN 1476-4687. doi: 10.1038/s41586-020-2649-2. URL https://www.nature.com/articles/s41586-020-2649-2.

47. B. Hassenstein and W. Reichardt. Systemtheoretische Analyse der Zeit-, Reihenfolgen- und Vorzeichenauswertung bei der Be-wegungsperzeption des Rüsselkäfers Chlorophanus. Zeitschrift für Naturforschung B, 11(9-10):513–524, Oct. 1956. ISSN 1865-7117, 0932-0776. doi: 10.1515/znb-1956-9-1004. URL https://www.degruyter.com/document/doi/10.1515/ znb-1956-9-1004/html.

48. S. He and W. R. Levick. Spatial-temporal response characteristics of the ON-OFF direction selective ganglion cells in the rabbit retina. Neuroscience Letters, 285(1):25–28, May 2000. ISSN 0304-3940. doi: 10.1016/S0304-3940(00)01030-2. URL https://www.sciencedirect.com/science/article/pii/S0304394000010302.

49. R. A. Holub and M. Morton-Gibson. Response of Visual Cortical Neurons of the cat to moving sinusoidal gratings: response-contrast functions and spatiotemporal interactions. Journal of Neurophysiology, 46(6):1244–1259, Dec. 1981. ISSN 0022-3077. doi: 10.1152/jn.1981.46.6.1244.

50. U. Homberg, R. A. Montague, and J. G. Hildebrand. Anatomy of antenno-cerebral pathways in the brain of the sphinx moth Manduca sexta. Cell and Tissue Research, 254(2):255–281, 1988. doi: 10.1007/BF00225800.

51. J. D. Hunter. Matplotlib: A 2D Graphics Environment. Computing in Science & Engineering, 9(3):90–95, May 2007. ISSN 1558-366X. doi: 10.1109/MCSE.2007.55. URL https://ieeexplore.ieee.org/document/4160265.

52. V. Jayaraman and G. Laurent. Evaluating a genetically encoded optical sensor of neural activity using electrophysiology in intact adult fruit flies. Frontiers in Neural Circuits, 1, Nov. 2007. ISSN 1662-5110. doi: 10.3389/neuro.04.003.2007. URL https://www.frontiersin.org/journals/neural-circuits/articles/10.3389/neuro.04.003.2007/full.

53. X. Jiang, E. Dimitriou, V. Grabe, R. Sun, H. Chang, Y. Zhang, J. Gershenzon, J. Rybak, B. S. Hansson, and S. Sachse. Ring-shaped odor coding in the antennal lobe of migratory locusts. Cell, 187(15):3973–3991.e24, July 2024. ISSN 0092-8674, 1097-4172. doi: 10.1016/j.cell.2024.05.036. URL https://www.cell.com/cell/abstract/S0092-8674(24)00580-4.

54. N. Kadakia, M. Demir, B. T. Michaelis, B. D. DeAngelis, M. A. Reidenbach, D. A. Clark, and T. Emonet. Odour motion sensing enhances navigation of complex plumes. Nature, 611(7937):754–761, Nov. 2022. ISSN 0028-0836, 1476-4687. doi: 10.1038/s41586-022-05423-4. URL https://www.nature.com/articles/s41586-022-05423-4.

55. N. D. Kathman, A. J. Lanz, J. D. Freed, and K. I. Nagel. Neural dynamics for working memory and evidence integration during olfactory navigation in Drosophila, Oct. 2024. URL https://www.biorxiv.org/content/10.1101/2024.10.05.616803v1. Pages: 2024.10.05.616803 Section: New Results.

56. J. S. Kennedy and D. Marsh. Pheromone-regulated anemotaxis in flying moths. Science, 184(4140):999–1001, May 1974. ISSN 0036-8075. doi: 10.1126/science.184.4140.999.

57. S. Kikuta, K. Sato, H. Kashiwadani, K. Tsunoda, T. Yamasoba, and K. Mori. Neurons in the anterior olfactory nucleus pars externa detect right or left localization of odor sources. Proceedings of the National Academy of Sciences, 107(27):12363–12368, July 2010. ISSN 0027-8424, 1091-6490. doi: 10.1073/pnas.1003999107. URL https://pnas.org/doi/full/10.1073/pnas.1003999107.

58. T. Kitamoto. Conditional modification of behavior in Drosophila by targeted expression of a temperature-sensitive shibire allele in defined neurons. Journal of Neurobiology, 47(2):81–92, May 2001. ISSN 0022-3034. doi: 10.1002/neu.1018.

59. N. C. Klapoetke, Y. Murata, S. S. Kim, S. R. Pulver, A. Birdsey-Benson, Y. K. Cho, T. K. Morimoto, A. S. Chuong, E. J. Carpenter, Z. Tian, J. Wang, Y. Xie, Z. Yan, Y. Zhang, B. Y. Chow, B. Surek, M. Melkonian, V. Jayaraman, M. Constantine-Paton, G. K.-S. Wong, and E. S. Boyden. Independent optical excitation of distinct neural populations. Nature Methods, 11(3):338–346, Mar. 2014. ISSN 1548-7091, 1548-7105. doi: 10.1038/nmeth.2836. URL https://www.nature.com/articles/nmeth.2836.

60. M. Knaden, A. Strutz, J. Ahsan, S. Sachse, and B. S. Hansson. Spatial representation of odorant valence in an insect brain. Cell Reports, 1(4):392–399, Apr. 2012. ISSN 2211-1247. doi: 10.1016/j.celrep.2012.03.002. URL https://www.sciencedirect.com/science/article/pii/S2211124712000733.

61. A. Kurtovic, A. Widmer, and B. J. Dickson. A single class of olfactory neurons mediates behavioural responses to a Drosophila sex pheromone. Nature, 446(7135):542–546, Mar. 2007. ISSN 1476-4687. doi: 10.1038/nature05672. URL https://www.nature.com/articles/nature05672.

62. S.-L. Lai and T. Lee. Genetic mosaic with dual binary transcriptional systems in Drosophila. Nature Neuroscience, 9(5):703–709, May 2006. ISSN 1546-1726. doi: 10.1038/nn1681. URL https://www.nature.com/articles/nn1681.

63. A. J. Lanz, N. D. Kathman, E. Hao, B. Ermentrout, and K. I. Nagel. Disinhibition of a recurrent attractor gates a persistent goal signal for navigation, Apr. 2026. URL https://www.biorxiv.org/content/10.1101/2025.10.07.681003v2. ISSN: 2692-8205 Pages: 2025.10.07.681003 Section: New Results.

64. N.-F. Liou, S.-H. Lin, Y.-J. Chen, K.-T. Tsai, C.-J. Yang, T.-Y. Lin, T.-H. Wu, H.-J. Lin, Y.-T. Chen, D. M. Gohl, M. Silies, and Y.-H. Chou. Diverse populations of local interneurons integrate into the Drosophila adult olfactory circuit. Nature Communications, 9 (1):2232, June 2018. ISSN 2041-1723. doi: 10.1038/s41467-018-04675-x. URL https://www.nature.com/articles/s41467-018-04675-x.

65. M. S. Livingstone and B. R. Conway. Substructure of Direction-Selective Receptive Fields in Macaque V1. Journal of Neurophysiology, 89(5):2743–2759, May 2003. ISSN 0022-3077, 1522-1598. doi: 10.1152/jn.00822.2002. URL https://www.physiology.org/doi/10.1152/jn.00822.2002.

66. A. Mafra-Neto and R. T. Cardé. Fine-scale structure of pheromone plumes modulates upwind orientation of flying moths. Nature, 369(6476):142–144, May 1994. ISSN 1476-4687. doi: 10.1038/369142a0. URL https://www.nature.com/articles/369142a0.

67. O. Mano, M. S. Creamer, C. A. Matulis, E. Salazar-Gatzimas, J. Chen, J. A. Zavatone-Veth, and D. A. Clark. Using slow frame rate imaging to extract fast receptive fields. Nature Communications, 10(1):4979, Oct. 2019. ISSN 2041-1723. doi: 10.1038/s41467-019-12974-0. URL https://www.nature.com/articles/s41467-019-12974-0. Publisher: Nature Publishing Group.

68. A. M. M. Matheson, A. J. Lanz, A. M. Medina, A. M. Licata, T. A. Currier, M. H. Syed, and K. I. Nagel. A neural circuit for wind-guided olfactory navigation. Nature Communications, 13(1):4613, Aug. 2022. ISSN 2041-1723. doi: 10.1038/s41467-022-32247-7. URL https://www.nature.com/articles/s41467-022-32247-7.

69. C. A. Matulis, J. Chen, A. D. Gonzalez-Suarez, R. Behnia, and D. A. Clark. Heterogeneous Temporal Contrast Adaptation in Drosophila Direction-Selective Circuits. Current Biology, 30(2):222–236.e6, Jan. 2020. ISSN 09609822. doi: 10.1016/j.cub.2019.11.077. URL https://linkinghub.elsevier.com/retrieve/pii/S0960982219315799.

70. W. McKinney. Data Structures for Statistical Computing in Python. In Data Structures for Statistical Computing in Python, pages 56–61, Austin, Texas, 2010. doi: 10.25080/Majora-92bf1922-00a. URL https://doi.curvenote.com/10.25080/ Majora-92bf1922-00a.

71. P. A. Moore, N. Scholz, and J. Atema. Chemical orientation of lobsters, homarus americanus, in turbulent odor plumes. Journal of Chemical Ecology, 17(7):1293–1307, July 1991. ISSN 0098-0331. doi: 10.1007/BF00983763.

72. J. Murlis, J. S. Elkinton, and R. T. Cardé. Odor Plumes and How Insects Use Them. Annual Review of Entomology, 37(Volume 37, 1992):505–532, Jan. 1992. ISSN 0066-4170, 1545-4487. doi: 10.1146/annurev.en.37.010192.002445. URL https://www.annualreviews.org/content/journals/10.1146/annurev.en.37.010192.002445.

73. D. Münch and C. G. Galizia. DoOR 2.0 - Comprehensive Mapping of Drosophila melanogaster Odorant Responses. Scientific Reports, 6(1):21841, Feb. 2016. ISSN 2045-2322. doi: 10.1038/srep21841. URL https://www.nature.com/articles/srep21841.

74. S. Namiki and R. Kanzaki. Comparative neuroanatomy of the lateral accessory lobe in the insect brain. Frontiers in Physiology, 7: 244, 2016. doi: 10.3389/fphys.2016.00244.

75. S. Namiki, S. Iwabuchi, P. Pansopha Kono, and R. Kanzaki. Information flow through neural circuits for pheromone orientation. Nature Communications, 5:5919, 2014. doi: 10.1038/ncomms6919.

76. A. Pauli, F. Althoff, R. A. Oliveira, S. Heidmann, O. Schuldiner, C. F. Lehner, B. J. Dickson, and K. Nasmyth. Cell-Type-Specific TEV Protease Cleavage Reveals Cohesin Functions in Drosophila Neurons. Developmental Cell, 14(2):239–251, Feb. 2008. ISSN 1534-5807. doi: 10.1016/j.devcel.2007.12.009. URL https://www.cell.com/developmental-cell/abstract/S1534-5807(07)00486-8.

77. T. A. Pologruto, B. L. Sabatini, and K. Svoboda. ScanImage: Flexible software for operating laser scanning microscopes. BioMedical Engineering OnLine, 2(1):13, May 2003. ISSN 1475-925X. doi: 10.1186/1475-925X-2-13. URL https://biomedical-engineering-online.biomedcentral.com/articles/10.1186/1475-925X-2-13.

78. J. Porter, B. Craven, R. M. Khan, S.-J. Chang, I. Kang, B. Judkewitz, J. Volpe, G. Settles, and N. Sobel. Mechanisms of scent-tracking in humans. Nature Neuroscience, 10(1):27–29, Jan. 2007. ISSN 1546-1726. doi: 10.1038/nn1819. URL https://www.nature.com/articles/nn1819.

79. M. Potters and W. Bialek. Statistical mechanics and visual signal processing. Journal de Physique I, 4(11):1755–1775, Nov. 1994. ISSN 1155-4304, 1286-4862. doi: 10.1051/jp1:1994219. URL 10.1051/jp1:1994219.

80. N. J. Priebe, S. G. Lisberger, and J. A. Movshon. Tuning for spatiotemporal frequency and speed in directionally selective neurons of macaque striate cortex. The Journal of Neuroscience: The Official Journal of the Society for Neuroscience, 26(11):2941–2950, Mar. 2006. ISSN 1529-2401. doi: 10.1523/JNEUROSCI.3936-05.2006.

81. T. P. Quinn and A. H. Dittman. Pacific salmon migrations and homing: mechanisms and adaptive significance. Trends in Ecology & Evolution, 5(6):174–177, June 1990. ISSN 0169-5347. doi: 10.1016/0169-5347(90)90205-R.

82. R. Rajan, J. P. Clement, and U. S. Bhalla. Rats Smell in Stereo. Science, 311(5761):666–670, Feb. 2006. ISSN 0036-8075, 1095-9203. doi: 10.1126/science.1122096. URL https://www.science.org/doi/10.1126/science.1122096.

83. W. Reichardt and A.-k. Guo. Elementary pattern discrimination (behavioural experiments with the fly Musca domestica). Biological Cybernetics, 53(5):285–306, Mar. 1986. ISSN 1432-0770. doi: 10.1007/BF00336562. URL 10.1007/BF00336562.

84. E. Salazar-Gatzimas, J. Chen, M. S. Creamer, O. Mano, H. B. Mandel, C. A. Matulis, J. Pottackal, and D. A. Clark. Direct Measurement of Correlation Responses in Drosophila Elementary Motion Detectors Reveals Fast Timescale Tuning. Neuron, 92(1):227–239, Oct. 2016. ISSN 1097-4199. doi: 10.1016/j.neuron.2016.09.017.

85. F. Salman, J. Jonaitis, J. D. Ralston, O. M. Cook, M. M. Bennett, T. R. Sizemore, C. J. Guarniere, K. L. Ramachandra, K. E. Coates, J. L. Fox, and A. M. Dacks. Connectivity of serotonin neurons reveals a constrained inhibitory subnetwork within the olfactory system. Journal of Neurophysiology, 136(1):77–93, July 2026. ISSN 0022-3077, 1522-1598. doi: 10.1152/jn.00571.2025. URL https://journals.physiology.org/doi/10.1152/jn.00571.2025.

86. J.-C. Sandoz and R. Menzel. Side-Specificity of Olfactory Learning in the Honeybee: Generalization between Odors and Sides. Learning & Memory, 8(5):286–294, Sept. 2001. ISSN 1072-0502, 1549-5485. doi: 10.1101/lm.41401. URL http://learnmem.cshlp.org/lookup/doi/10.1101/lm.41401.

87. L. K. Scheffer, C. S. Xu, M. Januszewski, Z. Lu, S.-y. Takemura, K. J. Hayworth, G. B. Huang, K. Shinomiya, J. Maitlin-Shepard, S. Berg, J. Clements, P. M. Hubbard, W. T. Katz, L. Umayam, T. Zhao, D. Ackerman, T. Blakely, J. Bogovic, T. Dolafi, D. Kainmueller, T. Kawase, K. A. Khairy, L. Leavitt, P. H. Li, L. Lindsey, N. Neubarth, D. J. Olbris, H. Otsuna, E. T. Trautman, M. Ito, A. S. Bates, J. Goldammer, T. Wolff, R. Svirskas, P. Schlegel, E. Neace, C. J. Knecht, C. X. Alvarado, D. A. Bailey, S. Ballinger, J. A. Borycz, B. S. Canino, N. Cheatham, M. Cook, M. Dreher, O. Duclos, B. Eubanks, K. Fairbanks, S. Finley, N. Forknall, A. Francis, G. P. Hopkins, E. M. Joyce, S. Kim, N. A. Kirk, J. Kovalyak, S. A. Lauchie, A. Lohff, C. Maldonado, E. A. Manley, S. McLin, C. Mooney, M. Ndama, O. Ogundeyi, N. Okeoma, C. Ordish, N. Padilla, C. M. Patrick, T. Paterson, E. E. Phillips, E. M. Phillips, N. Rampally, C. Ribeiro, M. K. Robertson, J. T. Rymer, S. M. Ryan, M. Sammons, A. K. Scott, A. L. Scott, A. Shinomiya, C. Smith, K. Smith, N. L. Smith, M. A. Sobeski, A. Suleiman, J. Swift, S. Takemura, I. Talebi, D. Tarnogorska, E. Tenshaw, T. Tokhi, J. J. Walsh, T. Yang, J. A. Horne, F. Li, R. Parekh, P. K. Rivlin, V. Jayaraman, M. Costa, G. S. Jefferis, K. Ito, S. Saalfeld, R. George, I. A. Meinertzhagen, G. M. Rubin, H. F. Hess, V. Jain, and S. M. Plaza. A connectome and analysis of the adult Drosophila central brain. eLife, 9: e57443, Sept. 2020. ISSN 2050-084X. doi: 10.7554/eLife.57443. URL https://elifesciences.org/articles/57443.

88. P. Schlegel, A. S. Bates, T. Stürner, S. R. Jagannathan, N. Drummond, J. Hsu, L. Serratosa Capdevila, A. Javier, E. C. Marin, A. Barth-Maron, I. F. Tamimi, F. Li, G. M. Rubin, S. M. Plaza, M. Costa, and G. S. X. E. Jefferis. Information flow, cell types and stereotypy in a full olfactory connectome. eLife, 10:e66018, May 2021. ISSN 2050-084X. doi: 10.7554/eLife.66018. URL https://elifesciences.org/articles/66018.

89. P. Schlegel, Y. Yin, A. S. Bates, S. Dorkenwald, K. Eichler, P. Brooks, D. S. Han, M. Gkantia, M. dos Santos, E. J. Munnelly, G. Badala-mente, L. Serratosa Capdevila, V. A. Sane, A. M. C. Fragniere, L. Kiassat, M. W. Pleijzier, T. Stürner, I. F. M. Tamimi, C. R. Dunne, I. Salgarella, A. Javier, S. Fang, E. Perlman, T. Kazimiers, S. R. Jagannathan, A. Matsliah, A. R. Sterling, S.-c. Yu, C. E. McKellar, M. Costa, H. S. Seung, M. Murthy, V. Hartenstein, D. D. Bock, and G. S. X. E. Jefferis. Whole-brain annotation and multi-connectome cell typing of Drosophila. Nature, 634(8032):139–152, Oct. 2024. ISSN 1476-4687. doi: 10.1038/s41586-024-07686-5. URL https://www.nature.com/articles/s41586-024-07686-5.

90. J. L. Semmelhack and J. W. Wang. Select Drosophila glomeruli mediate innate olfactory attraction and aversion. Nature, 459(7244): 218–223, Apr. 2009. ISSN 0028-0836, 1476-4687. doi: 10.1038/nature07983. URL https://www.nature.com/articles/nature07983.

91. P. Singh, S. Goyal, S. Gupta, S. Garg, A. Tiwari, V. Rajput, A. S. Bates, A. K. Gupta, and N. Gupta. Combinatorial encoding of odors in the mosquito antennal lobe. Nature Communications, 14(1):3539, June 2023. ISSN 2041-1723. doi: 10.1038/s41467-023-39303-w. URL https://www.nature.com/articles/s41467-023-39303-w.

92. M. F. Strube-Bloss, M. P. Nawrot, and R. Menzel. Neural correlates of side-specific odour memory in mushroom body output neurons. Proceedings of the Royal Society B: Biological Sciences, 283(1844):20161270, Dec. 2016. ISSN 0962-8452. doi: 10.1098/rspb.2016.1270. URL 10.1098/rspb.2016.1270.

93. H. Suzuki. Convergence of olfactory inputs from both antennae in the brain of the honeybee. Journal of Experimental Biology, 62(1): 11–26, 1975. doi: 10.1242/jeb.62.1.11.

94. I. Taisz, E. Donà, D. Münch, S. N. Bailey, B. J. Morris, K. I. Meechan, K. M. Stevens, I. Varela-Martínez, M. Gkantia, P. Schlegel, C. Ribeiro, G. S. Jefferis, and D. S. Galili. Generating parallel representations of position and identity in the olfactory system. Cell, 186(12):2556–2573.e22, June 2023. ISSN 00928674. doi: 10.1016/j.cell.2023.04.038. URL https://linkinghub.elsevier.com/retrieve/pii/S0092867423004725.

95. T. Takasaki, S. Namiki, and R. Kanzaki. Use of bilateral information to determine the walking direction during orientation to a pheromone source in the silkmoth Bombyx mori. Journal of Comparative Physiology A, 198(4):295–307, 2012. doi: 10.1007/s00359-011-0708-8.

96. R. Tanaka and D. A. Clark. Neural mechanisms to exploit positional geometry for collision avoidance. Current biology: CB, 32(11): 2357–2374.e6, June 2022. ISSN 1879-0445. doi: 10.1016/j.cub.2022.04.023.

97. D. Task, C.-C. Lin, A. Vulpe, A. Afify, S. Ballou, M. Brbic, P. Schlegel, J. Raji, G. S. Jefferis, H. Li, K. Menuz, and C. J. Potter. Chemoreceptor co-expression in Drosophila melanogaster olfactory neurons. eLife, 11:e72599, Apr. 2022. ISSN 2050-084X. doi: 10.7554/eLife.72599. URL 10.7554/eLife.72599.

98. L. Tirian and B. J. Dickson. The VT GAL4, LexA, and split-GAL4 driver line collections for targeted expression in the Drosophila nervous system, Oct. 2017. URL https://www.biorxiv.org/content/10.1101/198648v1. Pages: 198648 Section: New Results.

99. N. J. Vickers and T. C. Baker. Reiterative responses to single strands of odor promote sustained upwind flight and odor source location by moths. Proceedings of the National Academy of Sciences, 91(13):5756–5760, June 1994.10.1073/pnas.91.13.5756. URL https://www.pnas.org/doi/10.1073/pnas.91.13.5756.

100. P. Virtanen, R. Gommers, T. E. Oliphant, M. Haberland, T. Reddy, D. Cournapeau, E. Burovski, P. Peterson, W. Weckesser, J. Bright, S. J. van der Walt, M. Brett, J. Wilson, K. J. Millman, N. Mayorov, A. R. J. Nelson, E. Jones, R. Kern, E. Larson, C. J. Carey, . Polat, Y. Feng, E. W. Moore, J. VanderPlas, D. Laxalde, J. Perktold, R. Cimrman, I. Henriksen, E. A. Quintero, C. R. Harris, A. M. Archibald, A. H. Ribeiro, F. Pedregosa, and P. van Mulbregt. SciPy 1.0: fundamental algorithms for scientific computing in Python. Nature Methods, 17(3):261–272, Mar. 2020. ISSN 1548-7105. doi: 10.1038/s41592-019-0686-2. URL https://www.nature.com/articles/s41592-019-0686-2.

101. J. W. Wang, A. M. Wong, J. Flores, L. B. Vosshall, and R. Axel. Two-Photon Calcium Imaging Reveals an Odor-Evoked Map of Activity in the Fly Brain. Cell, 112(2):271–282, Jan. 2003. ISSN 0092-8674, 1097-4172. doi: 10.1016/S0092-8674(03)00004-7. URL https://www.cell.com/cell/abstract/S0092-8674(03)00004-7.

102. W. Wei. Neural Mechanisms of Motion Processing in the Mammalian Retina. Annual Review of Vision Science, 4(1):165–192, Sept. 2018. ISSN 2374-4642, 2374-4650. doi: 10.1146/annurev-vision-091517-034048. URL https://www.annualreviews.org/ doi/10.1146/annurev-vision-091517-034048.

103. R. I. Wilson, G. C. Turner, and G. Laurent. Transformation of Olfactory Representations in the Drosophila Antennal Lobe. Science, 303(5656):366–370, Jan. 2004. doi: 10.1126/science.1090782. URL https://www.science.org/doi/10.1126/science.1090782. Publisher: American Association for the Advancement of Science.

104. H. H. Yang and T. R. Clandinin. Elementary Motion Detection in *Drosophila*: Algorithms and Mechanisms. Annual Review of Vision Science, 4(1):143–163, Sept. 2018. doi: 10.1146/annurev-vision-091517-034153. URL https://www.ncbi.nlm.nih.gov/pmc/articles/PMC8097889/. Publisher: NIH Public Access.

105. D. M. Zimmerman, B. L. d. Bivort, and A. D. T. Samuel. The larval Drosophila mushroom body balances lateralized sensing and interhemispheric integration, Feb. 2026. URL https://www.biorxiv.org/content/10.1101/2025.03.02.641007v3. ISSN: 2692-8205 Pages: 2025.03.02.641007 Section: New Results.

